# Stress-Induced Mucin 13 Reductions Drive Intestinal Microbiome Shifts and Despair Behaviors

**DOI:** 10.1101/2022.10.13.512135

**Authors:** Courtney R. Rivet-Noor, Andrea R. Merchak, Caroline Render, Naudia Gay, Rebecca M. Beiter, Ryan Brown, Austin Keeler, G. Brett Moreau, Sihan Li, Deniz E. Olgun, Alexandra D. Steigmeyer, Rachel Ofer, Tobey Mihn Huu Phan, Kiranmayi Vemuri, Lei Chen, Keira E. Mahoney, Jung-Bum Shin, Stacy A. Malaker, Chris Deppmann, Michael Verzi, Alban Gaultier

**Affiliations:** Center for Brain Immunology and Glia, University of Virginia School of Medicine, Charlottesville, VA 22908, USA; Department of Neuroscience, University of Virginia School of Medicine, Charlottesville, VA 22908, USA; Graduate Program in Neuroscience, University of Virginia School of Medicine, Charlottesville, VA 22908, USA; Undergraduate Department of Global Studies, University of Virginia College of Arts and Sciences, Charlottesville, VA, 22904, USA; Department of Biology, University of Virginia College of Arts and Sciences, Charlottesville, VA, 22904, USA; Division of Infectious Diseases and International Health, Department of Medicine, University of Virginia School of Medicine, Charlottesville, Virginia, USA; Graduate Program in Biochemistry and Molecular Genetics, University of Virginia School of Medicine, Charlottesville, VA 22908, USA; Undergraduate Department of Computer Science, University of Virginia School of Engineering and Applied Science, Charlottesville, VA 22904, USA; Undergraduate Department of Neuroscience Studies, University of Virginia College of Arts and Sciences, Charlottesville, VA, 22904, USA; Department of Chemistry, Yale University, New Haven, CT 06511, USA; Department of Genetics, Human Genetics Institute of New Jersey, Rutgers Cancer Institute of New Jersey, Rutgers Center for Lipid Research, Division of Environmental & Population Health Biosciences, EOHSI, New Brunswick, New Jersey, 08901, USA; School of Life Science and Technology, Key Laboratory of Developmental Genes and Human Disease, Southeast University, Nanjing, China

## Abstract

Depression is a prevalent psychological condition with limited treatment options. While its etiology is multifactorial, both chronic stress and changes in the microbiome are associated with disease pathology. In depression, stress is known to induce microbiome dysbiosis, defined here as a change in microbial composition associated with a pathological condition. This state of dysbiosis is then known to feedback on depressive symptoms. While studies have demonstrated that targeted restoration of the microbiome can alleviate depressive-like symptoms in mice, translating these findings to human patients has proven challenging due to the complexity of the human microbiome. As such, there is an urgent need to identify factors upstream of microbial dysbiosis. Here we investigate the role of mucin 13 as an upstream mediator of microbiome composition changes. Using a model of chronic stress, we show that the mucosal protein, mucin 13, is selectively reduced after psychological stress exposure. We further demonstrate that the reduction of *Muc13* is mediated by the *Hnf4* transcription factor family. Finally, we determine that deleting *Muc13* is sufficient to drive microbiome shifts and despair behaviors. These findings shed light on the mechanisms behind stress-induced microbial changes and reveal a regulator of mucin 13 expression.

**Summary:** In this paper, authors identified a pathway by which stress induces microbiome shifts. They found that psychological stress selectively alters a key mucosal protein, mucin 13, which in turn modifies the microbial niche to induce changes in bacterial composition.

## Introduction

Depression and anxiety impact millions of people worldwide.^1^ Although many treatments exist for these disorders, high rates of treatment-resistant cases remain, with up to 60% of patients unable to find satisfactory results.^2^ This large population of patients living without relief for their mental health conditions highlights the need for further investigation into novel therapeutic treatments. While the etiology of depression remains complex, stress is known to be one of the largest contributing factors.^3,4^ In addition, stress and depression are both associated with changes in the gut microbiome.^5,6^ Alterations in the gut microbiome composition are present in depressed patients and conserved in animal models of depression, allowing researchers to investigate the bacterial strains that are most associated with disease onset. The microbiome has been extensively targeted as a potential treatment for mental health disorders: modification of the microbiome has been shown to reduce depression symptoms in humans and depressive-like behaviors in mice.^7–10^ While promising, efforts to broadly manipulate the human microbiome remain inconsistent due to the microbiome’s complexity, unknown microbe-microbe interactions, therapeutic microbes failing to colonize, microbe resource availability, and host heterogeneity.^9,11,12^ To circumvent current microbiome therapeutic limitations and increase chances of successful microbial alterations in patients, there is a critical need to identify key conserved, targetable regulators of the microbiome.

The mucosal layer is critical to maintaining both intestinal homeostasis and overall health. In the absence of mucosal proteins, known as mucins, sweeping microbiome changes and spontaneous disease occur.^13–17^ Mucins are highly glycosylated proteins that fall into two categories: soluble mucins and transmembrane mucins. ^14,18^ While soluble mucins make up the gel-like layer that is commonly associated with mucus, transmembrane mucins remain tethered to the cell membrane.^14,19^ In the intestines, mucin 2 is the dominant soluble mucin, while mucin 13 and mucin 17 make up the majority of the intestinal transmembrane mucins (collectively known as the glycocalyx).^14,19^ Both soluble and transmembrane mucins have important roles in regulating the intestine -- and importantly, the microbiome.^14^ The mucosal layer provides both a food source and anchor point for intestinal bacteria.^14^ In addition, recent works have demonstrated that the mucosal layer drives the stability and composition of the microbiome by providing a permissive state for bacterial growth, and selecting for specific bacterial types along the intestines by varying availability of glycan chains.^20–23^ Supporting this idea, deletion of core 1 O-glycans in mice results in dramatic alterations to the microbiome composition and spontaneous colitis.^24,25^

In addition, it has been demonstrated that stress hormones can alter the glycosylation patterns of mucins, supporting the hypothesis that stress induces alterations in the mucosal layer in a way that changes the microbial niche.^24,26,27^ Together, these studies provide a foundation for the hypothesis that stress can alter the mucus layer, and that this alteration can induce changes in microbial composition.

Given the strong evidence supporting the role of the mucosal layer in microbiome homeostasis and regulation, as well as the known ways in which stress can alter the mucosal layer, we hypothesized that exposure to chronic stress alters the mucosal layer to change the microbial niche and initiate microbiome changes. Here, we demonstrate that a model of chronic stress selectively alters the expression of a transmembrane mucin, mucin 13. This change is not seen after transfer of a microbiome from “stressed” mice into antibiotic treated or germ-free mice, suggesting that changes in the microbiome composition itself does not reduce mucin 13. In addition, we demonstrate the transcription factor hepatocyte nuclear factor 4 (HNF4) regulates *Muc13* expression by binding the *Muc13* promoter. Finally, we demonstrate that deletion of *Muc13* induces baseline despair behaviors, renders animals more susceptible to anxiety-like behaviors, and alters microbial signatures in a way that mimics the composition of microbes after stress exposure. Together, these findings suggest that Mucin 13 and HNF4 are mechanistic regulators of stress-induced microbiome shifts.

## Results

### Unpredictable Chronic Mild Restraint Stress Drives Despair and Anxiety-Like Behaviors and Modifies the Gut Microbiome Composition

To investigate the mechanisms of stress-induced microbiome alterations, we used unpredictable chronic mild restraint stress (UCMRS), a version of a model of chronic stress known to induce anxiety- and depressive-like behaviors and alter gut flora (Fig. 1A).^7^ After three weeks, mice exposed to UCMRS showed a significant increase in anxiety-like behaviors characterized by an increase in nestlet shredding and decrease in time spent in the open arms of the elevated plus maze (Fig. 1B). Despair behaviors were also detected through an increase in time inactive in both the tail suspension and forced swim tests. In addition, a reduction in sucrose preference, a measure for anhedonia, was observed in UCMRS exposed mice (Fig. 1B). In addition to behavioral readouts, levels of murine stress-associated markers were measured from serum by mass spectrometry (Fig. 1C and Ext. Data 1A-G). An increase in murine cortisol, but not corticosterone, was detected in stressed mice, confirming a robust chronic stress response (Fig. 1C Ext. Data 1A). While murine stress responses are often associated with changes in corticosterone, mice also express cortisol to a lesser extent.^28–30^ In addition, while changes in corticosterone are abundant in acute models of stress, others have suggested that cortisol levels remain elevated in chronic stress while corticosterone levels return to baseline.^28^ Additionally, decreases in both serotonin and glutamate were observed, consistent with findings in human depression.^31,32^ No other molecules tested were changed (Ext. Data 1A-G). This data highlights the validity of our model by demonstrating that changes in molecules associated with human anxiety and depression, also occur in mice exposed to UCMRS.

**Figure 1:**
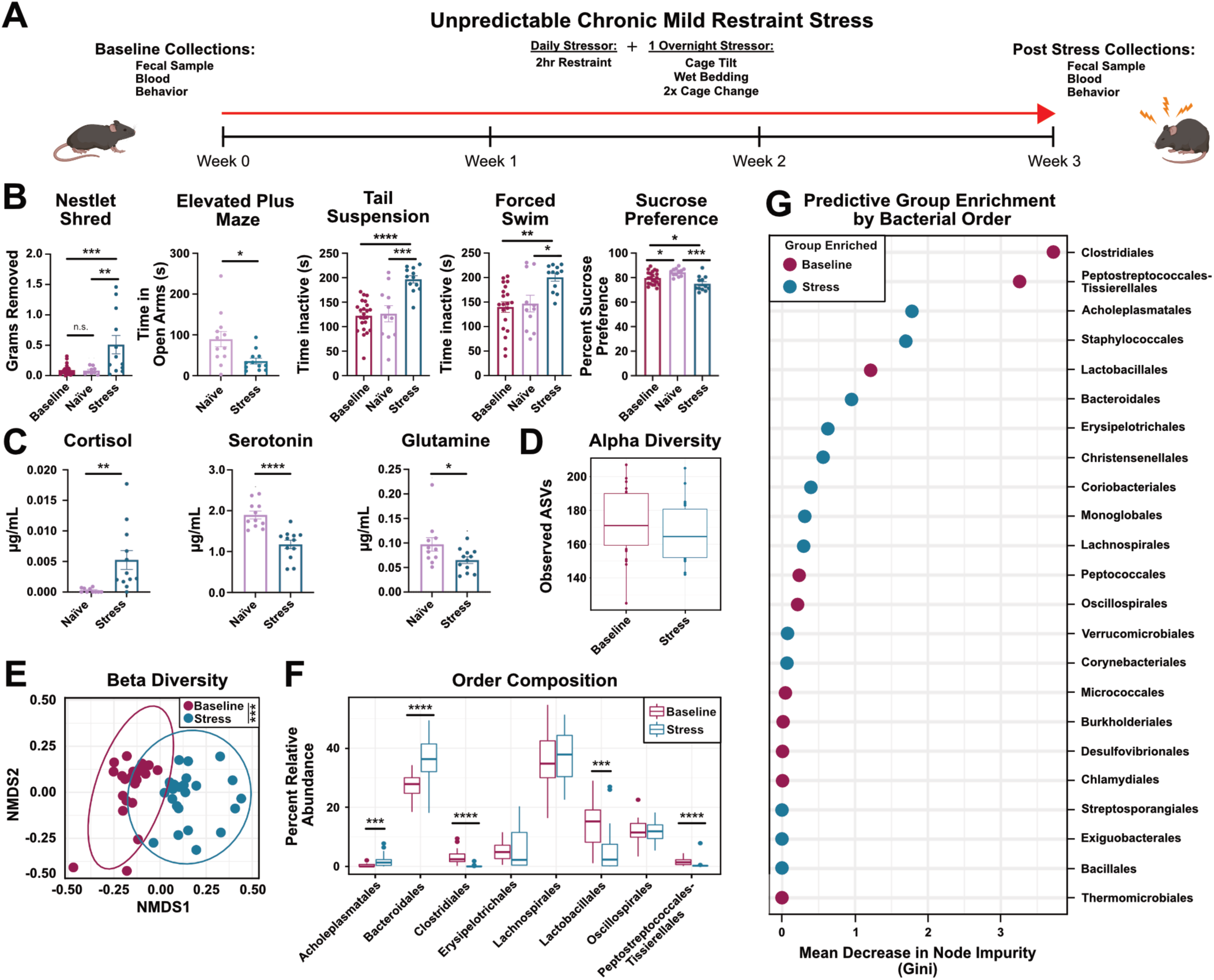
Unpredictable Chronic Mild Restraint Stress Induces Despair and Anxiety-Like Behaviors and Microbiome Dysbiosis: (A) Experimental Design. (B) Nestlet shred (One-way ANOVA), elevated plus maze (t-test), tail suspension (One-way ANOVA), forced swim (One-way ANOVA), and sucrose preference (One-way ANOVA) tests between baseline, naïve controls, and stress animals. Male mice, n=11/12 per group, representative of N=2. (C) Targeted mass spectrometry from serum of naïve or stress animals (t-tests, n=11/12 per group). (D) Alpha diversity plot showing microbial richness (observed ASVs) between baseline and stress mice (Wilcoxon Rank Sum test). (E) NMDS plot of Bray-Curtis dissimilarity between baseline and stress fecal microbiome samples (PERMANOVA). (F) Relative abundances of bacterial orders >1% (Wilcoxon Rank Sum test with Bonferroni correction). (G) Importance plot from a random forest model predicting bacterial orders that discriminate between baseline and stress groups. Importance is based on the Gini index where larger values are more important to model. Male mice, n=24/group, N=1.

We next evaluated stress-induced microbiome composition changes by performing 16S sequencing on fecal DNA from stressed and naïve animals. Using ASV-level data, we observed no differences in measures of alpha diversity (richness (Fig. 1D) or evenness (Ext. Data 1H)). However, there were significant differences in ASV-level derived beta diversity between naïve and stressed animals (Fig. 1E). These differences were driven by changes in community composition at the ASV level; for clarity, bacterial orders are depicted. Specifically, stress drove relative reductions in *Clostridiales*, *Lactobacillales*, and an expansion of *Bacteroidales* (Fig. 1F and Ext. Data 1I). Specific genus level changes between groups and the associated statistics can be found in Ext. Data 1J and Ext. Data 5, Tab 2. Lastly, an ASV based random forest model accurately predicted the bacterial orders associated with either baseline or stressed groups, strengthening the link between stress and specific microbial shifts (Fig. 1G). Taken together, these results demonstrate that unpredictable chronic mild restraint stress (UCMRS) induces despair and anxiety-like behaviors in mice, alters murine stress hormone levels, and induces composition shifts in the murine microbiome.

### Unpredictable Chronic Mild Restraint Stress Reduces *Muc13* Expression *In Vivo*

While it is well accepted that stress can change the microbiome in humans and mice (Fig. 1), the mechanisms behind these microbial shifts remain unknown.^9,33–36^ Because the mucosal layer, primarily composed of heavily glycosylated mucins, provides both an anchor point and nutrient reservoir for bacteria, we hypothesized that a change in mucin composition could induce microbiome shifts.^18,19,37^ To test this, we first evaluated if mucin expression changed in stressed conditions. Using RNA extracted from individual sections of the intestines of mice exposed to UCMRS and naïve controls (Fig. 2A), we examined mucin expression by qPCR. Of all the mucins examined (all detectable mucins: *Muc1*, *2*, *4*, *13*, and *17*), we found that *Muc13* was uniquely reduced across intestinal segments (Fig. 2B-C and Ext. Data 2A-C). Additionally, to examine mucin changes across genetic background, we examined *Muc13* and *Muc2* expression in BALB/cJ mice exposed to stress. We found a similar reduction in *Muc13*, while no changes in *Muc2* expression were observed (Ext. Data 2D-E). Taken together, these results suggest that stress induces changes in *Muc13* expression, and that these changes are conserved across different genetic backgrounds.

**Figure 2:**
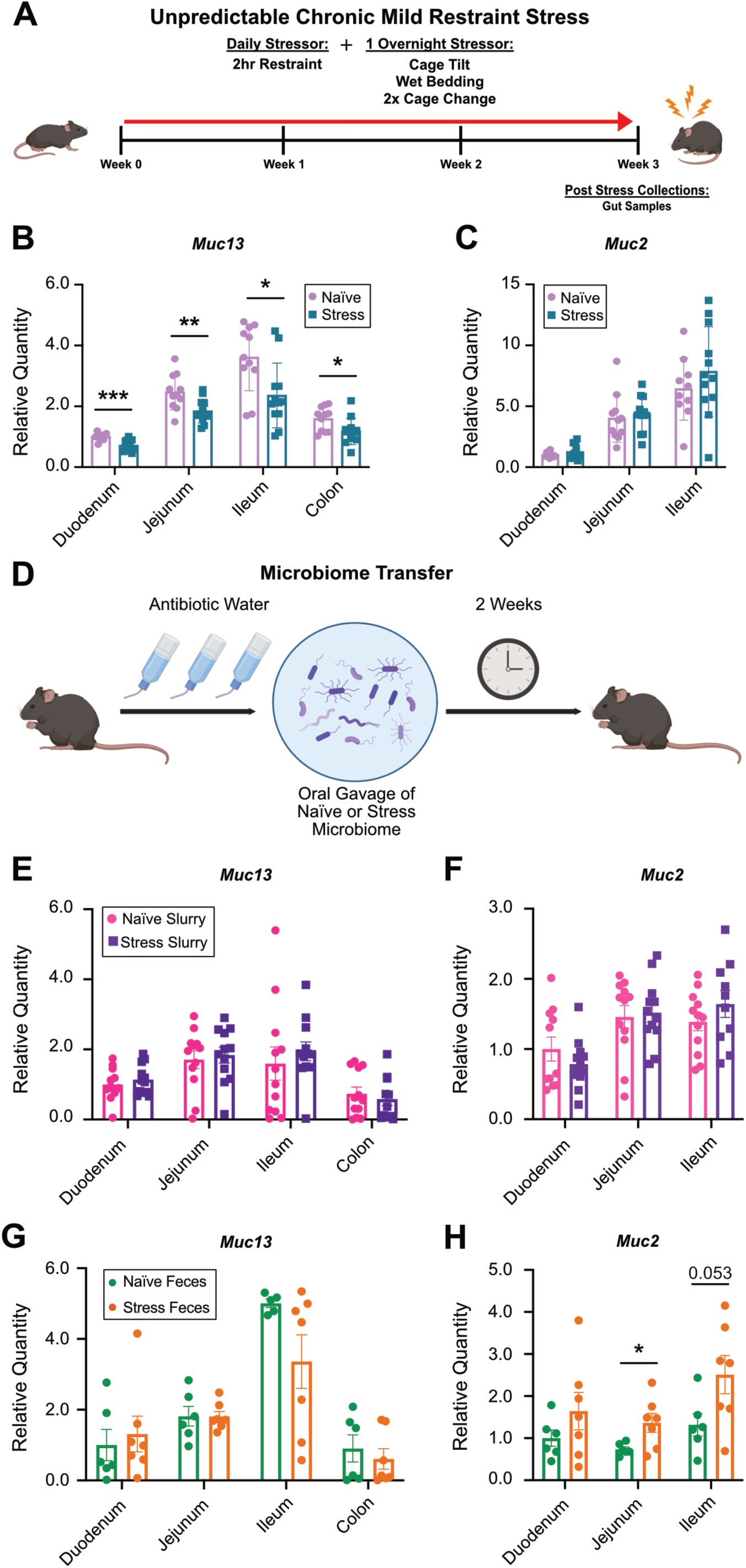
Stress Drives *Muc13* Reductions Independently of Transferrable Microbial Products: (A) Experimental design. Relative quantity of (B) *Muc13* and (C) *Muc2* transcripts by qPCR in individual sections of the intestine. *Muc2* expression in the colon was higher than housekeeping genes preventing quantification by qPCR. Unpaired two-tailed, t-tests. Male mice, n=11/12 per group, N=2. (D) Experimental design. Relative quantity of (E) *Muc13* and (F) *Muc2* transcripts in the intestines of animals receiving a naïve or stress fecal microbiome after antibiotic treatment via gavage. Male mice, n=12 per group. T-tests, two-tailed. Relative quantity of (G) *Muc13* and (H) *Muc2* in the intestines of germ-free mice given naïve or stress feces. Male mice, n=7 per group. T-tests, two-tailed.

### Stress-Induced *Muc13* Reductions are Driven Independently of Transferrable Microbial Products

The microbiome is a known influencer of the mucus layer. While we hypothesized that stress-induced changes in the mucosal layer lead to changes in microbiome composition, we also considered that stress-induced shifts in the microbiome could instead be modulating *Muc13* expression.^37,38^ If this was the case, it would suggest that mucin 13 changes would occur after microbial shifts in our model. To test this hypothesis, we utilized two different models to examine the impacts of the microbiome on mucin 13. First, after treating mice with antibiotics for two weeks, we transferred the fecal microbiomes from naïve or stressed mice via oral gavage (Fig. 2D). Two weeks post reconstitution, intestinal samples were collected to examine microbiome composition and mucin expression. 16S sequencing revealed that donor samples clustered distinctly from each other (Ext. Data 2F, light blue and dark pink). In addition, recipient mice clustered nearer to their donor sample but separately from each other, demonstrating distinct microbiomes and successful transfer of the expected signatures (Ext. Data 2F-G). We next examined RNA expression of intestinal mucins between groups. Importantly, no change in mucin 13 expression was detected between animals that received a stressed or naïve microbiome (Fig. 2E). In addition, we saw no change in any other mucins expressed between groups (Fig. 2F and Ext. Data 2H-J). These data demonstrate that microbial signatures from the transferred stressed microbiomes are not sufficient to reduce mucin 13 expression.

To complement this approach, we also examined the impact of stressed microbial signatures in germ free mice. Our lab has successfully demonstrated that microbial signatures from stressed mice can alter despair and anxiety-like behaviors in germ free mice.^39^ Therefore, using intestinal samples collected from germ free mice which were given fecal pellets from naïve or stress-exposed animals, we again examined changes in mucin expression by qPCR. Our results indicated that, similar to the antibiotic treated mice, no reduction in *Muc13* was observed (Fig. 2G). While we saw no changes in *Muc1*, *Muc4*, or *Muc17*, we did see an expected increase in *Muc*2 expression with microbiome reconstitution in germ free mice (Fig. 2H and Ext. Data 2K-M).^40^ These results demonstrate that microbial signatures from stressed animals, while enough to induce behavioral changes, are not sufficient to reduce *Muc13* expression in the intestines. This again suggests that the mucin 13 changes observed in stress are not driven by -- and indeed likely precede -- microbial changes in the intestines. Taken together, these results support the hypothesis that stress-induced reductions in mucin 13 are driven independently of transferrable microbial signatures.

### *Muc13* Reductions are Driven by Reductions in Intestinal HNF4

To understand how stress induces reductions in *Muc13* expression, we explored transcription factors which are known to bind to *Muc13* by examining published small intestine and colon ChIP-seq data sets through the UCSC genome browser.^41^ Results revealed that hepatocyte nuclear factor 4 (HNF4) is able to bind to the promoter region of *Muc13* in both the small and large intestines.^42,43^ HNF4, a transcription factor which is a member of the nuclear receptor superfamily, plays a critical role in regulating metabolism, epithelial tight junctions, and the differentiation and proliferation of enterocytes in the intestines.^44^ In addition to these functions, *Hnf*4 expression has been shown to regulate the brush border, the collection of microvilli along the intestinal tract.^45^ As the brush border is thought to be modulated by the density of the glycocalyx, and therefore transmembrane mucins, HNF4 is a logical candidate to mediate changes in *Muc13* expression.^46^ Importantly, stress hormones have also been shown to inhibit *Hnf*4 expression in a model of high fat diet.^47^ Together, these works provide a foundation for the hypothesis that stress may be able to regulate *Hnf4* expression in a way that reduces *Muc13* expression.

In order to investigate the ability of HNF4 to regulate *Muc13* expression, we first analyzed previously performed HNF4 and H3K27ac ChIP-seq on wildtype duodenal villus cells to examine binding at the *Muc1*3 locus.^48^ Results indicated that in wildtype mice, both HNF4 and H3K27ac, a marker of transcription, are enriched in the promoter region of *Muc13* (Fig. 3A). This suggests that HNF4 binds the *Muc13* promoter in a site that is marked for active transcription, supporting the idea that HNF4 can regulate *Muc13* expression. We next wanted to examine the impacts of HNF4 deletion on *Muc13* expression in the intestine. To do this, we utilized the previously described *Villin-Cre^ERT^*^2^; *Hnf4a*^F/F^; *Hnf4g* ^Crispr/Crispr^ double knockout mouse line (*Hnf4* DKO) which lacks both HNF4 paralogs.^48^ Analyzing previously performed RNA Polymerase II (RNAP2) ChIP-seq on duodenal villus epithelial cells in wildtype controls and *Hnf4* DKO animals, we found that the RNAP2 signal at the *Muc13* promoter is markedly reduced in the *Hnf4* DKO animals compared to controls (Fig. 3B). To complement this approach, we further examined previously performed H3K4me3 chromosome conformation capture with chromatin immunoprecipitation (HiChIP) and RNA-seq on control and *Hnf4* DKO duodenal villus epithelial cells.^49^ Analysis revealed fewer H3K4me3 chromatin loops in the *Muc13* promoter region in *Hnf4* DKO animals than in control duodenal villus epithelial cells, suggesting fewer sites of active transcription (Fig. 3C). In addition, RNA sequencing results showed a significant reduction in *Muc13* expression in duodenal villus epithelial cells after the loss of *Hnf4* (Fig. 3D). We next performed qPCR for mucin expression on all sections of the intestines from control and *Hnf4* DKO animals. We found that in addition to the successful deletion of the *Hnf4* paralogs (Fig. 3E-F), there was also a significant reduction in *Muc13* expression across the intestines (Fig. 3G), that was not seen in any other tested mucins (Fig 3H and Ext. Data 3A). Lastly, we analyzed intestinal expression of *Hnf4* from mice exposed to 3 weeks of UCMRS (Fig. 1A). Our results indicated that *Hnf4* expression is significantly reduced in the intestines after stress exposure (Fig. 3I-J). In addition, leucine rich repeat containing G protein coupled receptor 5 (Lgr5), which is known to be suppressed in *Hnf4* knockout animals, was also significantly reduced (Fig. 3K).^50^ Furthermore, germ-free mice that had been given fecal pellets from stressed or naïve mice showed no differences in *Hnf4* expression, suggesting that *Hnf4* expression is not driven by transferrable microbial products (Ext. Data 3B-C).

**Figure 3:**
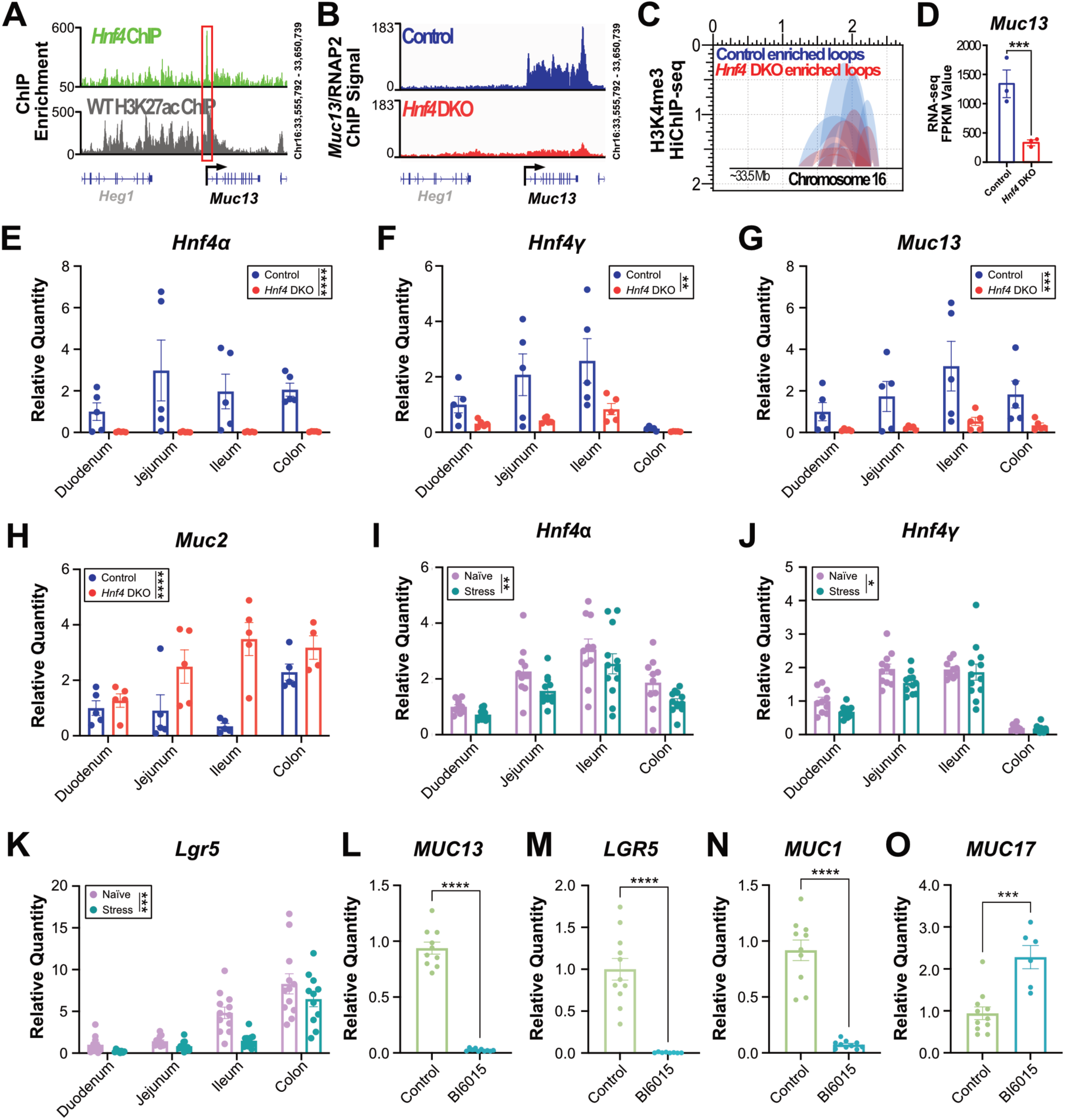
HNF4 Regulates Expression of *Muc13* both *In Vivo* and *In Vitro*: (A) HNF4 and H3K27ac ChIP-Seq of wildtype duodenal tissue at the *Muc13* promoter. (B) RNA Polymerase II ChIP-Seq in duodenal villus tissue in control and *Hnf4* DKO animals at the *Muc13* promoter. (C) H3K4me3 HiChIP-Seq examining chromatin loops in control and *Hnf4* DKO duodenal villus tissue at the *Muc13* promoter. (D) *Muc13* RNA-Seq fragments per kilobase per million in control and *Hnf4* DKO duodenal tissue (stats performed with embedded cuffdiff package). All sequencing experiments performed on 3 mice/group. Relative quantity of (E) *Hnf4α*, (F) *Hnf4γ*, (G) *Muc13*, and (H) *Muc2* across the intestines in control and *Hnf4* DKO animals. Female mice, 5 mice/group. Two-way ANOVA. Relative quantities of (I) *Hnf4α*, (J) *Hnf4γ*, and (K) *Lgr5* across the intestines in naïve and stress animals. Male mice, 11-12 mice per group; Two-way ANOVA. Relative quantities of (L) *MUC13*, (M) *LGR5*, (N) *MUC1*, and (O) *MUC17* in human HT-29 cells treated for 24 hours with 10μM of HNF4 antagonist BI6015 or DMSO control. N=3, n=11. Two-tailed unpaired t-tests.

In order to examine the impacts of HNF4 on *Muc13* expression more directly, we utilized HT-29 cells. HT-29 cells are a human colon cancer cell line that is known to express both mucins and *Hnf4*.^51,52^ Cells were treated with an HNF4 antagonist (BI6015) or a DMSO control for 24 hours.^53,54^ We then extracted the RNA from the collected cells and examined mucin expression as before. We found that treatment with the antagonist BI6015 showed a significant downregulation of *MUC13* expression (Fig. 3L). In addition, *LGR5* showed decreased expression with BI6015, demonstrating successful antagonist activity on HNF4 (Fig. 3M).^50^ Finally, BI6015 was also found to significantly decrease expression of *MUC1* and significantly increase expression of *MUC17* (Fig. 3N-O). However, as the expression of these other transmembrane mucins was not impacted by HNF4 reductions *in vivo* (Ext. Data 2B-C, Ext. Data 3A), there are likely other factors that more strongly contribute to changes in their expression. These results further support the hypothesis that *MUC13* is regulated by HNF4.

Together, these data suggest that in the absence of HNF4, there is less active transcription at, and fewer transcripts of, the *Muc13* gene; this supports the hypothesis that *Muc13* is regulated by HNF4. Our results also demonstrate that after UCMRS exposure, there is a significant reduction in *Hnf4* that occurs independently of transferrable microbial products, suggesting that stress is capable of reducing *Hnf4 in vivo*. Finally, we show that direct modulation of HNF4 results in *Muc13* expression changes *in vitro,* validating our *in vivo* findings. Taken together, these results demonstrate that *Muc13* expression is modulated by HNF4 both *in vitro* and *in vivo,* and after stress exposure.

### *Muc13* Deletion Induces Baseline Behavioral and Microbiome Changes and Renders Mice More Susceptible to UCMRS

Given our data demonstrating that *Muc13* is specifically downregulated in stressed animals (Fig. 2B), we sought to examine if deleting *Muc13* impacted the microbiome and behavioral outputs in mice. Using the *i*-GONAD system, we created a *Muc13* knockout (*Muc13*^-/-^) mouse line by deleting a 475bp region of the *Muc13* gene containing the start codon (Fig. 4A).^55,56^ Validation of the knockout line was performed using both qPCR (Fig. 4B-C) and immunofluorescence (Fig. 4D-E). Importantly, no changes were observed in *Muc2* at the transcript or protein levels, suggesting that *Muc2* expression was not changed with the loss of *Muc13* (Fig. 4C and F). To complement this approach, we also examined detectable mucins at the protein level by mass spectrometry in *Muc13^-/-^* and control animals. To do this, we employed a mucin enrichment strategy that takes advantage of an inactive protease selective for mucin domains; mucins from *Muc13^-/-^* and control animal intestines were enriched and subjected to MS analysis.^57,58^ We were able to see a significant reduction in the MUC13 protein in all sections of the intestine (Fig. 4G). No other detectable mucin was statistically significantly changed (Fold Change >5), suggesting that none of the other membrane mucins nor soluble mucins, compensate for the loss of MUC13. Together, these data demonstrate a successful deletion of MUC13 in our knockout line.

**Figure 4:**
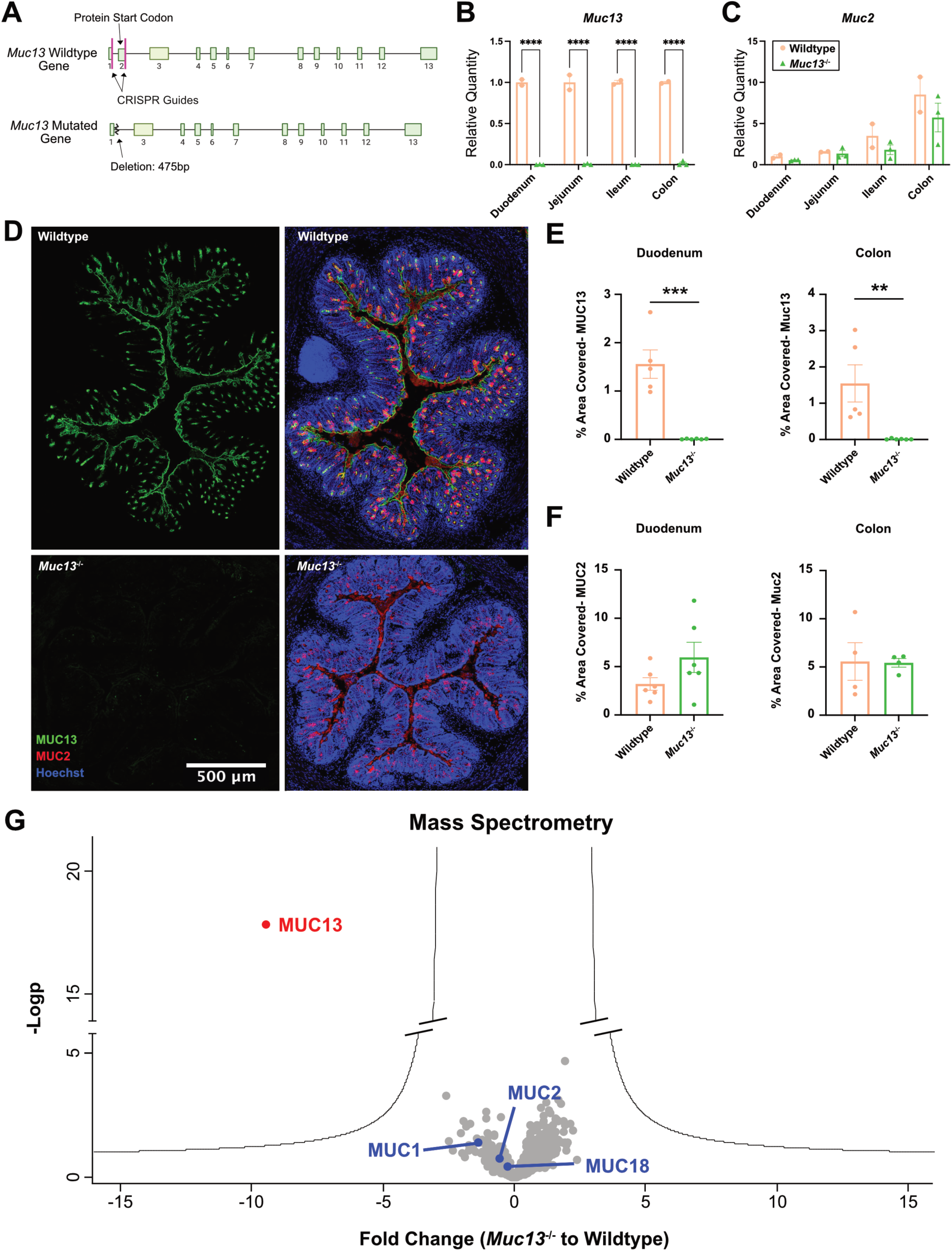
*Muc13* Deletion Validation: (A) *Muc13* deletion schematic. Relative quantities of (B) *Muc13* and (C) *Muc2* in wildtype and *Muc13^-/-^* mice. Male mice, n=2-3 per group, N=1. T-tests, two-tailed. (D) Representative images of immunofluorescence (IF) staining of MUC13, MUC2 and Hoechst in the colon of wildtype and *Muc13^-/-^* animals. Quantification of IF of (E) MUC13 and (F) MUC2 in the duodenum and colon of wildtype and *Muc13^-/-^* mice. Male mice. T-tests, n=4-6 per group, N=2. (G) Fold change of mucins in the intestines of *Muc13^-/-^* compared to wildtype controls by mass spectrometry. All sections of the intestines are pooled for a total of 12 samples from 3 mice per group. Male mice.

We next sought to understand the impacts of *Muc13* deletion on microbiome composition. We compared 16S sequencing results from fecal samples collected from adult wildtype and *Muc13*^-/-^ animals that were separated by genotype at weaning. Interestingly, we found that *Muc13*^-/-^ animals (light green) clustered distinctly from wildtype controls (light orange), suggesting that *Muc13* deletion significantly impacts the composition of the microbiome at baseline (Fig. 5B). In addition, to understand if *Muc13^-/-^* animals have stress induced microbiome shifts, we used an abbreviated 1-week UCMRS protocol (Fig. 5A). In our hands, this abbreviated protocol allows for substantial microbial shifts, but does not yet yield all stress-induced anxiety-like and despair behaviors in wildtype animals (Fig. 5D-G, light orange to dark pink bars). This allows us to examine if *Muc13^-/-^* animals are susceptible to different UCMRS induced microbial shifts than wildtype controls, and if *Muc13^-/-^* mice are more susceptible to UCMRS-induced behavioral changes. Our results show that while wildtype samples show a significant shift in microbial composition after 1 week of UCMRS (light orange to dark pink dots), *Muc13*^-/-^ (light green to dark green dots) samples remained clustered together (Fig. 5B). This suggests that the microbiome of *Muc13*^-/-^ animals is different from WT animals at baseline and is less modified by 1-week of UCMRS exposure. In addition, samples from wildtype animals exposed to 1 week of UCMRS shifted towards both groups (baseline and 1 week UCMRS, light and dark green dots) of the *Muc13*^-/-^ animals, supporting the idea the *Muc13*^-/-^ animals have microbial signatures which mimic stress-induced microbiome signatures (Fig. 5B). To further understand the similarities between the *Muc13*^-/-^ microbiome and the UCMRS-exposed microbiome, we compared the significantly changed bacterial families between baseline wildtype and baseline *Muc13*^-/-^ animals to the significantly changed bacterial families between baseline wildtype animals and wildtype animals exposed to 1 week of UCMRS. Of the bacterial families that had significant changes in either group, 69% of those changes overlapped (Fig. 5C). While we also examined overlapping changes at the genus and ASV levels, algorithm matching limitations hindered our ability to draw conclusions. While 58% of genera and 62% of ASV changes overlapped between groups, only ∼45% of total reads could be matched to a known genus or ASV -- making clear comparisons at these levels difficult (Ext. Data 5, Tab 3). Nonetheless, data at the family level suggests significant overlap in changes between baseline control and baseline *Muc13^-/-^* animals when compared to baseline control and UCMRS exposed controls. Together, these data further suggest that *Muc13* deletion induces a change in the microbiome that closely mimics the microbial shifts observed after UCMRS exposure.

**Figure 5:**
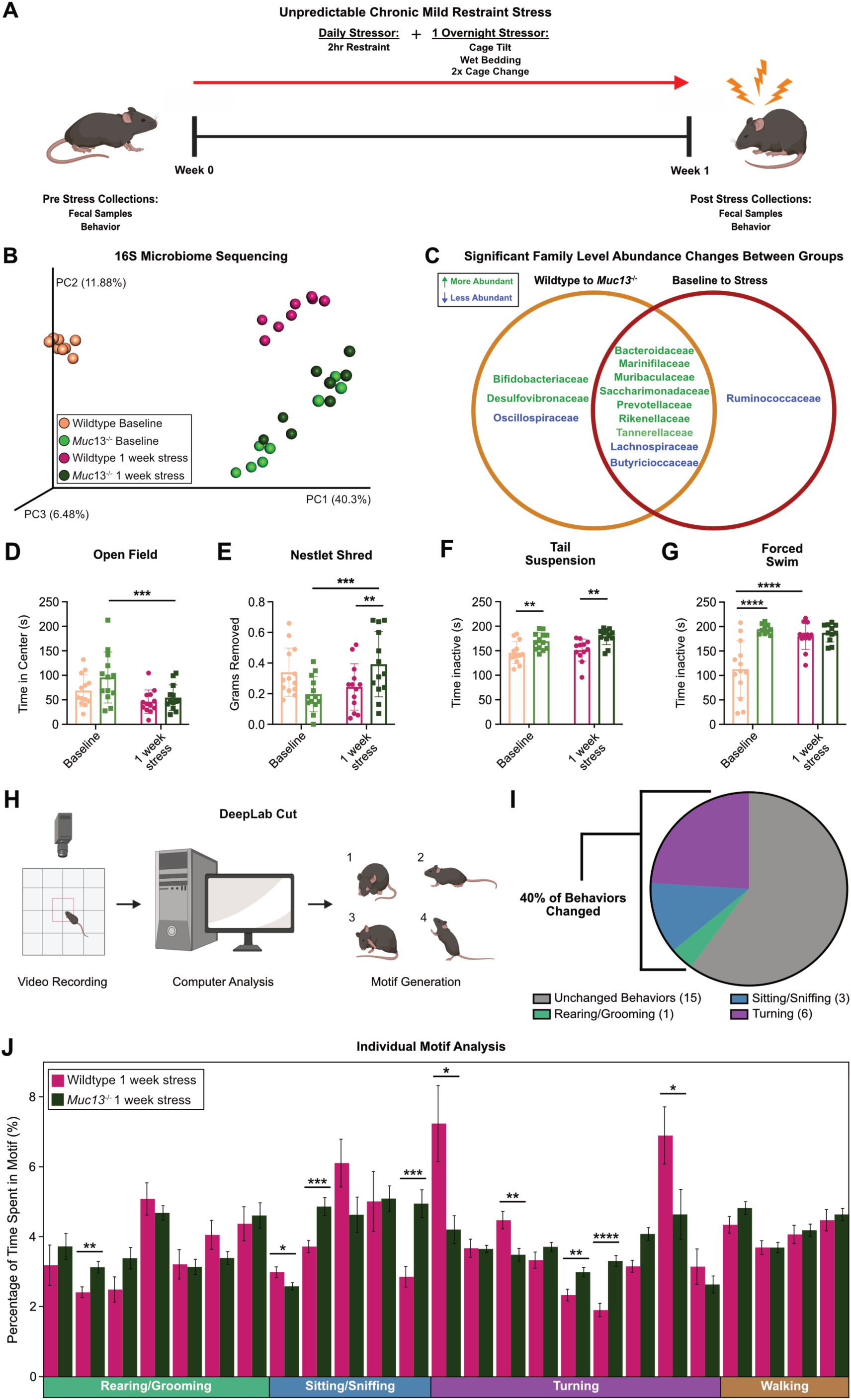
Microbiome and Behavioral Changes in *Muc13*^-/-^ Mice at Baseline and after Stress Exposure: (A) Experimental design. (B) PCoA plot of 16S fecal microbial sequencing in wildtype and *Muc13*^-/-^ animals at baseline and after 1 week of stress exposure. Male mice, n=8 per group, N=1. (C) Venn diagram comparing significantly changed families from 16S fecal microbiome sequencing between wildtype and *Muc13*^-/-^ animals to families changes between wildtype baseline and stress exposed animals. (D) Open field and (E) nestlet shredding tests comparing anxiety-like behaviors between wildtype and *Muc13*^-/-^ animals at baseline and after 1 week of stress exposure. (F) Tail suspension and (G) forced swim tests between wildtype and *Muc13*^-/-^ animals at baseline and after 1 week of stress exposure. Male mice, Two-way ANOVA, n=13 per group, N=2. (H) DeepLabCut experimental design. (I) Pie chart of quantified behavioral motifs changed between wildtype and *Muc13*^-/-^ animals. (J) Individual motif analysis after 1 week of stress exposure between wildtype and *Muc13*^-/-^ animals. Two-tailed T tests, male and female mice, n=13-24 per group, N=3.

Given the established connection between microbiome changes and mental health, we sought to examine if *Muc13* deletion impacted anxiety-like and despair behaviors in mice.^7^ Both *Muc13*^-/-^ animals and wildtype controls were subjected to the open field, nestlet shred, tail suspension, and forced swim tests (Fig. 5A, D-G). Behavior was collected at baseline and after 1 week of UCMRS for all animals. Interestingly, at baseline, we observed no differences in anxiety-like behaviors in the open field or nestlet shredding tests between *Muc13*^-/-^ and wildtype controls (Fig. 5D-E, left columns). However, strong despair behaviors were observed in both the tail suspension and forced swim tests (Fig. 5F-G, left columns), suggesting *Muc13*^-/-^ mice display innate despair, but not anxiety-like, behaviors. In addition, to gain a full picture of the impact of *Muc13* deletion on susceptibility to stress-induced despair and anxiety-like behaviors, we subjected wildtype and *Muc13*^-/-^ mice to 1 week of UCMRS. Critically, as supported by other studies, 1 week of UCMRS exposure represents a subclinical model of stress that does not induce all expected behavioral changes in wildtype animals^59,60^. Thus, we hypothesized that if *Muc13*^-/-^ deletion rendered animals more susceptible to stress, they would exhibit despair and anxiety-like behaviors earlier than wildtype animals.^61^ Supporting this hypothesis, following 1 week of UMCRS, *Muc13*^-/-^ animals had significant reductions in the amount of time spent in the center of the open field test, as well as significant increases in the amount of nestlet removed in the nestlet shred test. These results indicate that 1 week of UCMRS is sufficient to induce anxiety-like phenotypes in *Muc13*^-/-^ animals, but not their wildtype counterparts (Fig. 5D-E, green columns). Furthermore, in the tail suspension and forced swim tests, we saw no increase in time spent inactive as the *Muc13*^-/-^ animals presented with despair behaviors at baseline and likely hit a ceiling effect (Fig. 5F-G, green columns). To complement these classical behavioral assays, we used unbiased artificial intelligence (DeepLabCut) to detect animal poses (Fig. 5H-J and Ext. Data 4).^62,63^ In this approach, 10-minute videos of mice exploring an open field box were broken down into 25 individual motifs (Fig. 5H). Each motif was characterized and grouped into a general behavioral classification (rearing/grooming, turning, etc.). Individual motifs represent a behavioral pattern based on mouse limb and tail position detected by DeepLabCut. For example, the general behavior classification “turning” included motifs such “head turning left”, “mid-point of right turn”, etc. As these motifs all make up a complete behavior, there were grouped in general classifications. Once grouped, changes in the percentage of time spent in each behavioral motif were quantified between *Muc13*^-/-^ and wildtype animals (Fig. 5H-J). Baseline analysis revealed that of the 25 distinct motifs, 16% were significantly changed between groups (Ext. Data 4A-B), suggesting that, like in classical behavioral assays, distinctions between *Muc13*^-/-^ and wildtype animals are present at baseline. Similarly, the DeepLabCut software was able to detect significant differences between groups after 1 week of UCMRS exposure, with 40% of behavioral motifs being changed between *Muc13*^-/-^ and wildtype controls (Fig. 4I-J).

Our results show that *Muc13* deletion can drive both microbiome shifts that mirror microbial changes induced by stress, and despair behaviors at baseline -- highlighting the critical role of mucin 13 in maintaining microbial and behavioral homeostasis. In addition, deletion of *Muc13* renders animals more susceptible to stress-induced anxiety-like behaviors. Together, these results support the hypothesis that stress induced *Muc13* reductions can drive microbial shifts that lead to behavioral changes in mice.

## Discussion

Our work demonstrates that stress induces microbiome changes and behavioral changes in mice. In addition, stress also reduces expression of a key component of the mucosal layer, mucin 13. Our data also supports the hypothesis that stress mediates this reduction in *Muc13* expression by reducing the intestinal transcription factor HNF4. Furthermore, we demonstrate that deletion of mucin 13 is sufficient to induce microbiome shifts and behavioral changes that mimic stress-induced phenotypes at baseline. In addition to these baseline changes, mice lacking mucin 13 also are more susceptible to behavioral changes after exposure to a model of sub-clinical unpredictable chronic mild restraint stress. Together, these data define how stress is able to induce shifts in microbial composition in the intestines, providing a foundation for future studies.

Our results are supported by concepts in previously published literature. For example, it is well known that there is a reciprocal relationship between the mucus layer and microbes.^37^ In fact, disruption of the intestinal mucus layer results in sweeping microbiome changes, heightened inflammation, and disease onset--supporting the idea that changes in mucus can impact other systems in the body.^13,16,17,37,64–68^ Additionally, stress has been shown to alter the glycosylation patterns of mucins.^27^ As changes in glycosylation are also known to alter the microbiome, the connection between stress and mucin induced microbiome changes is well supported.^23,69^ Finally, the connection between mucins and depression has also been suggested in the literature. Specifically, single nucleotide polymorphisms (SNPs) in *MUC13* have been identified in GWAS studies of depressed populations.^70–72^ In addition, SNPs in O-glycosylation have also been identified in populations with treatment resistant depression.^72,73^ In aggregate, these data support our results, suggesting that mucin 13 is an important driver of microbiome shifts and despair behaviors in mice.

Mechanistically, our results show that HNF4 can bind to the *Muc13* promoter, while the loss of *Hnf4* results in reduced transcription of *Muc13* in the intestines of mice. In addition, we have shown that chronic stress induces reductions in both *Muc13* and *Hnf4* expression. While more work is needed to understand how stress regulates *Hnf4* in our chronic stress model, literature suggests that this regulation could occur through the phosphorylation of extracellular signal-regulated kinase (ERK) 1/2. While there is no glucocorticoid response element (GRE) in the *Hnf4* gene, ERK 1/2 is known to be a potent inhibitor of *Hnf4* expression.^74^ In addition, stress hormones have repeatedly been demonstrated to enhance ERK 1/2 phosphorylation.^75–77^ These works provide a mechanistic foundation for how stress could be reducing *Hnf4* expression in the intestines. While the basis for our hypothesis is well supported, no works have demonstrated how stress specifically regulates *Hnf4*, leaving a gap in knowledge to be filled by future studies.

By defining a novel mechanism through which stress alters the microbiome, our results have brought to light a key aspect in stress-induced behavioral changes. We have demonstrated that a transmembrane mucin is specifically and indirectly regulated by stress in a way that interferes with its homeostatic expression patterns, inducing microbiome composition changes and despair behaviors in mice. Mucin 13 is conserved between humans and mice, meaning that it could be a broadly applicable therapeutic target for patients with stress-induced depression, who present with microbiome changes. Although our results are directly related to stress-induced depression, they provide a basis for further research to target transmembrane mucins as intervention points for diseases that present with or are driven by pathogenic microbiome alterations, such as colitis or Parkinson’s Disease.^78^

## Methods

### Mice

C57BL/6j and BALB/cJ mice were purchased from Jackson Laboratories (strain #000664 and #000651, respectively). Mice were bred in-house and kept on a 12-h light/dark schedule. All mice were group housed up to 5 mice per cage unless separated for fighting or designated to a stressed group, where they were singly housed. All behavioral interventions were performed between 8 am and 3 pm and animals were sacrificed between 7 am and 1 pm. Experimental and control animals were sacrificed in alternating patterns to ensure proper controls for circadian consideration. Villin-Cre^ERT2^; Hnf4a^F/F^; Hnf4g^Crispr/Crispr^ animals were generated at Rutgers University under the care of the Verzi lab. Inducible knockouts for sequencing were given 1 injection of tamoxifen (50mg/kg) for 2-3 days (2-3 total doses) to catch early changes in the intestines. HNF4 DKO mice used for qPCR were given 1 injection of tamoxifen (50mg/kg) for 4 days (4 total doses). Animals were harvested on the 5^th^ day. All control animals were given saline.

Mucin-13 knockout lines were generated using the iGONAD technique as described, using a BTX ECM 830 Electroporation System (Harvard Apparatus).^55,56^ Briefly, the Muc13 sequence was taken from the UCSC genome browser, mouse assembly Dec. 2011 (GRCm38/mm10), Genomic Sequence (chr16:33,794,037-33,819,927)^41^. Exons 1 and 2 including the intervening intron where analyzed with CRISPOR (http://crispor.tefor.net/; Concordet and Haeussler 2018)^93^. Exon 2, containing the protein start sequence, was targeted for excision at these two target sequences + PAM sites: Exon2_protein_start_sequence GCAAGAGCAGCTACCATGAA (AGG) and Exon2_end_of_exon AGTCTCCTTTTGGTGACCGT (GGG). Alt-R S.p. HiFi Cas9 Nuclease, tracrRNA, and crRNA XT for the two target sequences were purchased (IDT). Prior to surgery, the Alt-R CRISPR/Cas9 reagents were prepared according to IDT guidelines: crRNA-XT and tracrRNA were annealed to form the gRNA, then complexed to the S.p HiFi Cas9 nuclease, and then diluted with sterile Opti-MEM with Fast Green FCF to aid visualization.

Plugged females were anesthetized and maintained with isoflurane ∼16 hours after copulation was assumed to have occurred (estimate copulation around midnight, surgeries were done at 4pm). Anesthesia was confirmed by toe pinch and eyes were lubricated with Puralube ointment. The lower dorsal skin of the mouse was soaked in betadine followed by two washes with isopropanol. A dorsal incision was made to expose the ovaries, oviduct, and uterine horn. Using a pulled micropipette, ∼1.5ul of the CRISPR/Cas9 solution was drawn up and then injected between the infundibulum and ampulla, intraoviductally dispensing the solution slowly to avoid backflow. The oviduct was covered in a wetted, sterile kimwipe and immediately electroporated using 3-mm platinum tweezer rode electrodes (BTX) delivering 50V for 5ms with 1s interval for 8 square-wave pulses using a BTX ECM 830 Electroporation System (Harvard Apparatus). The oviduct was returned to the abdominal cavity and the procedure was repeated on the opposite side oviduct. Once both oviducts were injected and electroporated the incision was sutured, and the stitches secured with liquid bandage before receiving a postop subcutaneous injection of ketoprofen (5mg/kg of mouse body weight) with subsequent injections for up to five days as needed. All procedures were approved by the University of Virginia ACUC (protocol #3918). All experiments were conducted and reported according to ARRIVE guidelines.

## METHOD DETAILS

### Stress Experiments and Behavioral Tests

Unpredictable Chronic Mild Restraint Stress (UCMRS) experiments and behavioral experiments (the forced swim, tail suspension, sucrose preference, open field, elevated plus maze, and nestlet shred tests) were performed as previously described.^62^

### Stress protocol

For UCMRS, all animals were single housed. Mice were exposed to 2 hours of restraint stress and one overnight stressor per day. Restraint stress was performed by putting mice into ventilated 50mL conical tubes for 2 hours. Overnight stressors included one of the following: 45-degree cage tilt, wet bedding, or 2x cage change. Wet bedding was performed by wetting standard cage bedding with ∼200mL of murine drinking water. 2x cage change included replacing cage bases 2x within 24 hours. 45-degree cage tilt included propping cages up to ∼45-degrees overnight.

### Behavioral Tests

All testing (with exception to the nestlet shred and sucrose preference tests) was recorded on a Hero Session 5 GoPro and analyzed with Noldus behavioral analysis software.

*Nestlet Shred:* Mice were moved into fresh individual cages and allowed to habituate for 1 hour. After 1 hour, a weighed nestlet was placed in the center of the cage. Mice were allowed to interact with the nestlet for 30 min. After 30 min, nestlets were removed and weighed. Before weighing, excess nestlet was stripped from the main piece of nestlet by lightly dragging fingers across each area of the nestlet starting from the center (center up, center left, center right, center down). Once completed, nestlet was flipped and the process was repeated for the corners (center to upper right, center to upper left, center to lower right, center to lower left). Equal pressure was applied to each nestlet. Change in amount shredded and percent shredded were calculated from the weighed nestlet values.

*Sucrose Preference test:* Mice were housed individually and given 2 water bottles/cage. One water bottle contained normal drinking water, the other contained 1% sucrose water. Mice had access to the bottles for 3 days. On Day 0, bottles were primed and weighed and put in cages. Day 1: 24 hours later, water bottles were removed from cages and weighed. Once weighed, bottles were replaced, but with swapped positions (i.e.-if sucrose was on the left on Day 0, it was on the right on Day 1). The first night of sucrose preference is considered habituation and the values are not included in analysis. Day 2: 24 hours after replacing water bottles, bottles were weighed and replaced as on day 1. Day 3: 24 hours after replacing water bottles, bottles were removed and weighed for a final time. Sucrose preference was calculated by determining the percentage of each bottle drunk on day 2 and 3 and averaging those values.

*Elevated Plus Maze:* Mice were placed on an elevated plus maze apparatus and allowed to explore for 5 minutes. The elevated plus maze was cleaned with 70% ethanol between each run and recorded on a GoPro Hero Session 5 or 8. Files were analyzed with Noldus behavior software.

*Open field/DeepLabCut:* Animals were placed into a 14in x 14in box with opaque walls and a raised clear bottom (18in). Animals were allowed to explore freely for 10 minutes. Hero session 8s were placed ∼18in below the box to record animal movements from below. Noldus behavioral software was used to determine if animals were in the perimeter or center of the box. Videos were also used for DeepLabCut analysis. Analysis is described below.

### Metabolite Mass Spectrometry

25 µL of plasma was extracted using 500 µL of cold acetonitrile followed by vortexing and 10 minutes of centrifugation at 14K rpm. 450 µL of the resulting supernatant was transferred to a clean Eppendorf and dried completely by Speedvac. Each sample was reconstituted with 25 µL of 50% methanol in 0.1% formic acid/water, vortexed and transferred to autosampler vials for analysis by mass spectrometry. Eleven metabolites were quantitated by PRM (parallel reaction monitoring) using a Thermo Orbitrap ID-X mass spectrometer coupled to a Waters BEH C18 column (15 cm x 2.1 mm). Metabolites were eluted with a Vanquish UHPLC system over a 15 min gradient (10 µL injection, 200 µL/min, buffer A –0.1% formic acid in water, buffer B - 0.1% formic acid in methanol). The mass resolution was set to 120K for detection using the transitions outlines in Extended Data 5, Tab 1.

Raw data files for samples, blanks and calibration curves were imported into Skyline (https://skyline.ms/project/home/begin.view). Calibration curves were generated with a linear regression. Peak areas for analytes in samples were used for quantification based in the generated calibration curves. Raw files can be downloaded for analysis at https://www.metabolomicsworkbench.org under study ID: ST002345

### Mass Spectrometry

Frozen intestinal tissue sections were homogenized use the Cryo-Cup Grinder (BioSpec Products, Cat. No. 206) according to manufacturer’s instructions. Pulverized tissue was transferred into ice-cold 150 uL dialyzable lysis buffer, consisting of 1% N-octylglucoside (Research Products International, N02007), 1% CHAPS (Research Products International, C41010), 50 mM Tris (American Bio, AB14044), 100 mM NaCl (Research Products International, S23025), 2 mM MgCl_2_ (American Bio, AB09006), benzonase (Sigma Aldrich, E1014), and protease inhibitor (Roche, 11836170001). Samples were rotated at 4 °C for 2 hours, followed by centrifugation for 30 minutes at 15,000 rcf and passed through a 0.7 µm filter. StcE^E447D^ was expressed and purified as previously described.^57,58^ 500 ug of NHS-Activated Agarose Slurry (Thermo Fisher Scientific, 26200) was activated with 1 mM HCl and PBS. StcE^E447D^ (1 mL of 1.45 mg/mL) was added to the slurry and rotated at 4 °C for 3 hours. Beads were rinsed with 100 mM Tris and the reaction quenched with 100 mM acetate buffer. Beads were washed with 20 mM Tris. Beads were then washed and stored in 20 mM Tris 150 mM NaCl. Filtered supernatants were rotated overnight at 4 °C with 100 uL StcE^E447D^-NHS bead slurry and 0.01 M EDTA. All reactions were brought up to 1 mL with 20 mM Tris 150 mM NaCl. Beads were then rinsed with 10 mM Tris 1 M NaCl, 10 mM Tris 150 mM NaCl, 10 mM NaCl and MS grade water (Thermo Scientific, 51140). Protein was eluted with 1% sodium deoxycholate (Research Products International, D91500-25.0) in 50 mM Ammonium bicarbonate (AmBic) (Honeywell Fluka, 40867), at 95 °C for 5 minutes. Dithiothreitol (DTT) (Sigma Aldrich, D0632) was added to 20% of the eluent to a final concentration of 2mM and allowed to react at 65 °C for 25 minutes. Alkylation in 5 mM iodoacetamide (IAA) (Sigma Aldrich, I1149) was performed for 15 minutes in the dark at room temperature. 0.05μg of sequencing-grade trypsin (Promega, V5111) was added to each 20% aliquot and allowed to react for 3 hours at 37 °C. Reactions were quenched by adding 1 µL formic acid. All reactions were diluted to a volume of 100 µL prior to desalting. Desalting was performed using 10mg Strata-X 33µm polymeric reversed phase SPE columns (Phenomenex, 8B-S100-AAK). Each column was activated using 1 mL acetonitrile (ACN) (Honeywell, LC015) followed by of 1 mL 0.1% formic acid, 1 mL 0.1% formic acid in 50% ACN, and equilibration with addition of 1 mL 0.1% formic acid. After equilibration, the samples were added to the column and rinsed with 200 µL 0.1% formic acid. The columns were transferred to a 1.5 mL tube for elution by two additions of 150 µL 0.1% formic acid in 50% ACN. The eluent was then dried using a vacuum concentrator (LabConco) prior to reconstitution in 10 µL of 0.1% formic acid.

Samples were analyzed by online nanoflow liquid chromatography-tandem mass spectrometry using an Orbitrap Eclipse Tribrid mass spectrometer (Thermo Fisher Scientific) coupled to a Dionex UltiMate 3000 HPLC (Thermo Fisher Scientific). For each analysis, 5μL was injected onto an Acclaim PepMap 100 column packed with 2 cm of 5µm C18 material (Thermo Fisher, 164564) using 0.1% formic acid in water (solvent A). Peptides were then separated on a 15cm PepMap RSLC EASY-Spray C18 Column packed with 2µm C18 material (Thermo Fisher, ES904) using a gradient from 0-35% solvent B. (0.1% formic acid with 80% CAN) in 60 minutes. Full scan MS1 spectra were collected at a resolution of 60,000, an automatic gain control of 1E5, and a mass range from 400-1500 m/z. Dynamic exclusion was enabled with a repeat count of 2 and repeat and duration of 8 seconds. Only charge states 2 to 6 were selected for fragmentation. MS2s were generated at top speed for 3 seconds. Higher-energy collisional dissociation (HCD) was performed on all selected precursor masses with the following parameters: isolation window of 2m/z, stepped collision energies of 25%, 30%, 40%, orbitrap detection (resolution of 7500), maximum inject time of 75ms, and a standard AGC target.

Label-free quantification was performed using the minimal workflow for MaxQuant according to the established protocol for standard data sets.^79^ Files were searched with fully specific cleavage C-terminal to an Arg or Lys residue, with 2 allowed missed cleavages. Carbamidomethyl Cys was set as a fixed modification. Raw data generated from each intestinal section was included and analyzed using Perseus, according to the recommended protocol for label-free interaction data (Fold change >5 as the cutoff).

### Fecal DNA Extraction and 16S rRNA Gene Sequencing

DNA was isolated from fecal pellets using the phenol/chloroform method as described.^7^ Samples were then processed with QIAquick PCR purification kit (Qiagen #28106). Full protocol is outlined below:

250μL of 0.1 mm zirconia/silica beads (BioSpec #11079101z) were added to a sterile 2 mL screwtop tube (Celltreat #230830). One 4 mm steel ball was added to each tube (BioSpec #11079132ss). 500μL of Buffer A (200 mM TrisHCl, pH 8.0, 200 mM NaCl, 20 mM EDTA) and 210μL of 20% SDS were added to each fecal pellet in a separate 1.7mL tube. Pellets were vortexed and supernatant transferred to the 2mL screwtop tube containing beads. 500μL of Phenol/chloroform/IAA were added to each screwtop tube. Tubes were allowed to beadbeat on high for 4 min at room temperature. Samples were then centrifuge at 12000 rpm for 5 min at 4°C. 420μL of the aqueous layer were transferred to a new 1.7mL tube. 100uL of samples were then processed using the Qiagen PCR Purification Kit

### 3 week UCMRS experiments

The V4 region of the 16S rRNA gene was amplified from each sample using a dual indexing sequencing strategy.^80^ Samples were sequenced on the MiSeq platform (Illumina) using the MiSeq Reagent Kit v2 (500 cycles, Illumina #MS102-2003) according to the manufacturer’s protocol with modifications found in the Schloss Wet Lab SOP (https://github.com/SchlossLab/MiSeq_WetLab_SOP).

### Fig 1 16S Sequence Analysis

All processing and analysis of 16S rRNA sequencing data was performed in R (version 4.1.2).^81^ Raw sequencing reads were processed for downstream analysis using DADA2 (version 1.22.0).^82^ Processing included inspection of raw reads for quality, filtering of low-quality reads, merging of paired reads, and removal of chimeric sequences. Length distribution of non-chimeric sequences was plotted to ensure lengths matched the expected V4 amplicon size. Taxonomy was assigned to amplicon sequence variants (ASVs) by aligning reads with the Silva reference database (version 138.1).^83^

Microbiota diversity and community composition were analyzed using the packages phyloseq (version 1.38.0), microbiome (version 1.16.0), and vegan (version 2.5.7).^84–86^ The packages tidyverse (version 1.3.0), and ggplot2 (version 3.3.5) were used for data organization and visualization.^87,88^ Random forest analysis was performed using the and randomForest (version 4.6.14), vegan (version 6.0.90), and pROC (version 1.18.0) packages.^89–91^

Samples were first divided into training (70% of samples, divided equally between Baseline and Stressed samples) and test (30% of samples) sets. The training set was used to tune the “mtry” parameter of the model, while the test set was used to validate model performance. Feature importance was determined using the Gini index, which measures the total decrease in node impurity averaged across all trees.

Custom code and detailed instructions can be found at: https://github.com/gbmoreau/Gautier-manuscript. Raw sequencing reads can be found in the NCBI Sequence Read Archive (SRA) database under PRJNA867333.

### Microbiome Transfer Experiments

Antibiotic microbiome transfer experiments were performed by treating mice with a cocktail of the following antibiotics: Ampicillin (Sigma-Aldrich, A8351), Neomycin (Sigma-Aldrich, N6386), Metronidazole (Sigma-Aldrich, M1547), and Vancomycin (Sigma-Aldrich, V1130). Ampicillin, Neomycin, and Metronidazole were added to water containing 8g/L of a zero-calorie sweetener at 1g/L. Vancomycin was added at 500mg/L. Mice were allowed to drink ad labium for two weeks and fresh antibiotic solutions were made once per week. After 2 weeks of antibiotic treatment, mice were given an oral gavage of 100uL of either naïve or stressed fecal slurry (equal weights of collected fecal pellets homogenized in 4mL PBS) every 2 days for a total of 4 gavage treatments. Animals were allowed to habituate to the introduced microbiome for 14 days from the first gavage. Germ free experiments were performed as described.^39^

### Microbiome Transfer and 1 week UCMRS 16S

Fecal pellet DNA extraction and 16S sequencing of the V4 region was performed by Zymo according to their protocols. Raw sequencing reads can be found in the NCBI Sequence Read Archive (SRA) database under PRJNA878703 and PRJNA866924, respectively.

### Cell Culture

HT-29 cells were purchased from ATCC and were grown in McCoy’s 5A media (ATCC 30-2007) as per the ATCC website. Cells were plated at a density of 150,000/well and allowed to adhere to 12 well plates overnight in complete media. Cells were treated with 10μM of either NTC (Sigma-Aldrich, SMB00208) or BI6015 (Sigma-Aldrich, 375240) for 24hrs. After 24 hours, media was aspirated, and cells were frozen at -80 until RNA extraction was performed.

### RNA and ChIP Seq Experiments

All ChIP Seq experiments were performed previously on duodenal villus tissue as described.^48,49^ Raw data can be found under GEO accession numbers GSE112946 (RNA-seq and ChIP-seq) and GSE148691 (HiChIP-seq).

### DeepLabCut

Deeplabcut analysis was performed as previously described.^62^ Full protocol is outlined below:

Animal pose estimation was performed by using a deep-learning package called DeepLabCut. The detailed mathematical model and network architecture for DeepLabCut was previously published by the Mathis lab ^63^. We generated a DeepLabCut convolutional neural network to analyze our open field test videos, which is trained in a supervised manner: 16 manually labeled points were selected as references of transfer learning, including nose, left and right eyes, left and right ears, neck, left and right arm, left and right leg, 3 points on body and 3 points on tail. In total, 15 randomly selected videos were used for building a training dataset. Finally, the performance of the neural network is evaluated by human researchers.

Estimated mouse poses from DeepLabCut were further analyzed by using a package called Variational Animal Motion Embedding (VAME), which classifies animal behavior in an unsupervised manner. VAME was developed by the Pavol Bauer group and the package can be downloaded at https://github.com/LINCellularNeuroscience/VAME92. We trained a unique VAME recursive neural network for each experiment, which classifies each frame of the open field test video into 1 of the 25 behavioral motifs. Then, all behavior motifs were annotated and evaluated by blinded human researchers. With annotated frames, we were able to calculate the percentage of time usage of each motif, which is then used for principal component analysis and Kullback-Leibler divergence analysis. Custom code is available upon request.

### RNA Extraction and Quantitative PCR

For RNA extraction, cultured cells were pelleted, frozen, and lysed. RNA was extracted using the Bioline Isolate II RNA mini kit as per manufacture’s protocol (BIO-52073). RNA was quantified with a Biotek Epoch Microplate Spectrophotometer. Normalized RNA was reverse transcribed to cDNA with either the Bioline SensiFast cDNA Synthesis Kit (BIO-65054) or Applied Sciences High-Capacity cDNA Reverse Transcriptase Kit (43-688-13). cDNA was amplified using the Bioline SensiFast NO-ROX kit (BIO-86020), according to manufacturer’s instructions. Results were analyzed with the relative quantity (ΔΔCq) method. Probes used are listed in Extended Data 5, Tab 4.

### Immunofluorescence Staining and Imaging

1cm sections of intestinal samples were harvested and snap frozen in an OCT block (Fisher Scientific #14-373-65). Samples were cut on a cryostat in 15μM sections and placed on microscope slides. Tissue was fixed directly on the slides in carnoy’s solution (60% Ethanol, 30% Chloroform, and 10% Glacial Acetic Acid) for 10 min at room temperature. Tissue was washed in 1x PBS for 5 min 3x at room temperature. Antigen retrieval was performed with Tris-EDTA by bringing solution to a boil (∼30sec) in a microwave. Slides were washed in 1x PBS 3x for 5 min at room temperature. After final wash, sections were outlined with a hydrophobic pen (Vector Laboratories #H-4000). Samples were blocked for 1 hour at room temperature with 1:200 FC block in TBST (1x TBS + 0.3% Triton X-100) plus 2% Normal Donkey Serum (Jackson ImmunoResearch #017-000-121). Supernatant was removed and samples were blocked with Mouse on Mouse IgG for 1 hour at room temperature (Vector Laboratories #MKB-2213-1) plus 2% Normal Donkey Serum. Supernatant was removed and primary antibodies were added. Samples were stained with 1:1000 Mucin 13 (G-10) (Santa Cruz Biotechnology #sc-390115) and 1:1000 Mucin 2 (GeneTex #GTX100664) overnight at 4 degrees Celsius in blocking solution (1x PBS, 0.5g bovine serum albumin, 0.5% Triton X-100) + 2% normal donkey serum. Control samples were stained in blocking solution + 2% normal donkey serum with mouse IgG (Jackson ImmunoResearch #015-000-003) or rabbit IgG (Jackson ImmunoResearch #011-000-003) diluted to match the primary antibodies. The next day samples were washed in TBST for 5 min 3x at room temperature. Samples were then incubated in secondary antibodies for 3 hours at room temperature in blocking solution + 2% normal donkey serum. Secondary antibodies used were Donkey Anti-Mouse 488 (Jackson ImmunoResearch #715-545-150) and Donkey Anti-Rabbit 647 (Jackson ImmunoResearch #711-605-152). Secondaries were added at 2mg/mL. Samples were then stained with Hoechst (1:700) (Thermo Fisher Scientific #H3570) for 15 min in blocking solution + 2% normal donkey serum. Samples were washed 2x with TBST for 5 min at room temperature. Samples were washed a final time in TBS for 5 min at room temperature. Finally, slides imaged on a Leica Stellaris 5 confocal microscope.

## QUANTIFICATION AND STATISTICAL ANALYSIS

### Statistical analysis

All statistical analyses-except those associated with DeepLabCut, Mass Spectrometry, and 16S were performed in GraphPad Prism 9. Analyses comparing two groups were performed using a two-tailed T test. If the variances between groups were significantly different, a Welch’s correction was applied. Outliers were excluded if they fell more than two standard deviations from the mean. For all analyses, the threshold for significance was at p < 0.05. Repeats for each experiment are specified in the figure legend corresponding to the respective panel. Additional statistical detail, including all p-values, can be found in Ext. Data 5, Tab 5.

## Acknowledgements

The mass spectrometry experiments in Figure 1 were performed in the UVA Biomolecular Research Facility Core. The W.M. Keck Biomedical Mass Spectrometry Laboratory is funded by a grant from the University of Virginia’s School of Medicine. Alexandria Battison provided additional technical assistance in Mass Spectrometry experiments.

## Contributions

C.R.N. and A.G. designed the study; C.R.N., A.R.M., C.R., N.M.G., R.M.B., R.B., A.K., R.O., A.D.S., K.E.M., S.A.M., M.V, and C.D. performed and conceptualized experiments; G.B.M., S.L., D.E.O, T.M.H.P., K.V., L.C., and J.B.S. performed sequencing and DeepLabCut analysis; C.R.N and A.G. analyzed and wrote the manuscript; A.G. oversaw the project.

## Competing Interests

S.A.M. is an inventor on a Stanford patent related to the use of mucinases as research tools; she is also a consultant for InterVenn Biosciences.

## Funding

The authors are supported by grants from the UVA Trans University Microbiome Initiative pilot grant, Cat Miller Foundation, and the Owens Family Foundation. C.R.N is supported by the UVA Presidential Fellowship in Neuroscience. A.R.M. is supported by the NINDS T32 NS115657 and F31 AI174782. Work from the R.O., K.V., and L.C. from the lab of M.V. was supported by NIH grants R01DK121915 and R01DK126446. A.D.S. is supported by the National Institutes of Health Chemical Biology Training Grant (T32 GM067543). K.E.M. is supported by the Yale Endowed Postdoctoral Fellowship in the Biological Sciences.

## EXTENDED DATA

**Extended Data 1:**
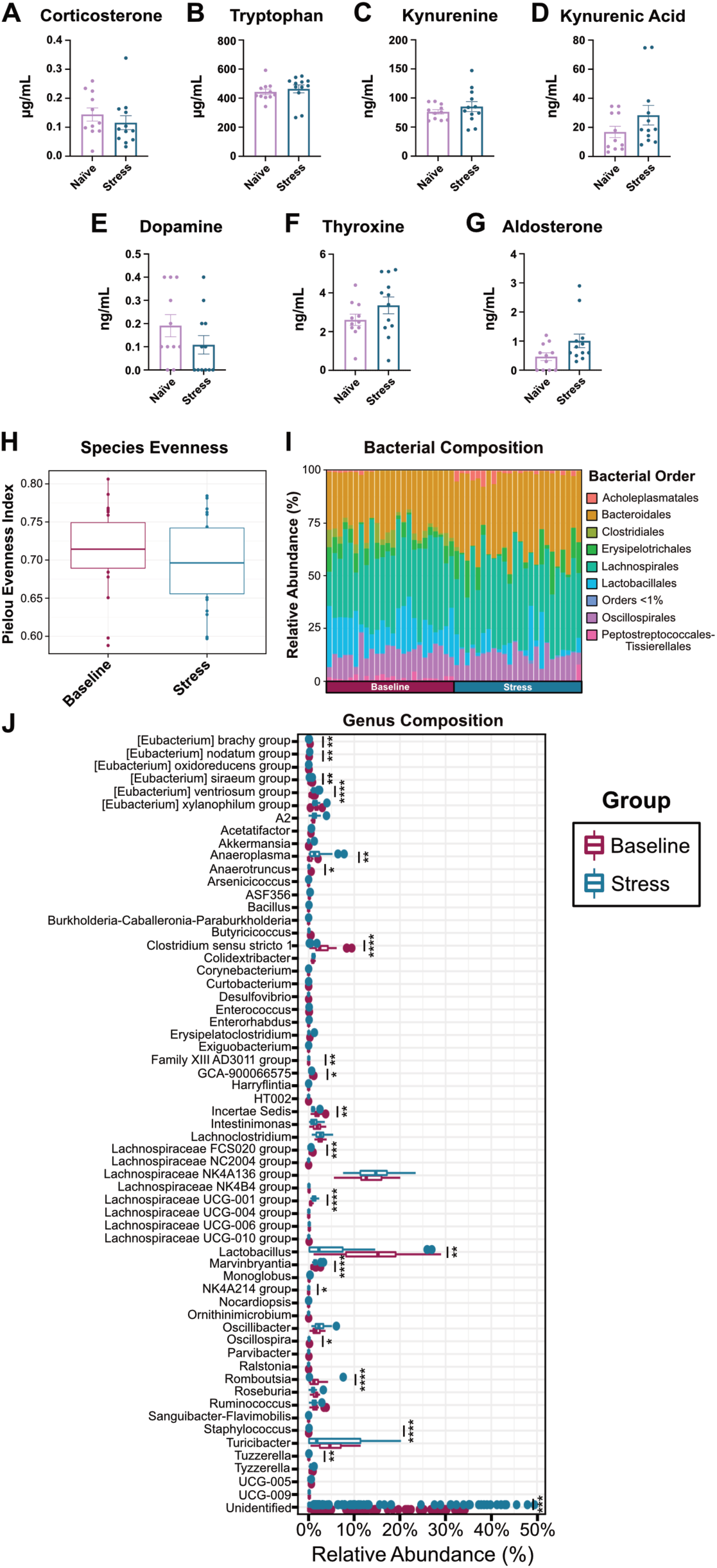
Serum Metabolites and 16S Sequencing after Stress Exposure: Mass spectrometry analysis of serum from naïve and stressed mice. (A) corticosterone, (B) tryptophan, (C) kynurenine, (D) kynurenic acid, (E) dopamine, (F) thyroxine, and (G) aldosterone. Male mice, n=11/12 per group, N=1, Unpaired two-tailed T-tests. MS Transitions can be found in Ext. Data5, Tab 1. (H) Pielou’s evenness index for baseline and stress mice. (I) Individual order relative abundances for baseline and stress mice. (J) Relative Abundances of the genus level between naïve and stress mice. Male mice, n=24/group, N=1. P-values for (J) can be found in Ext. Data 5, Tab 2.

**Extended Data 2:**
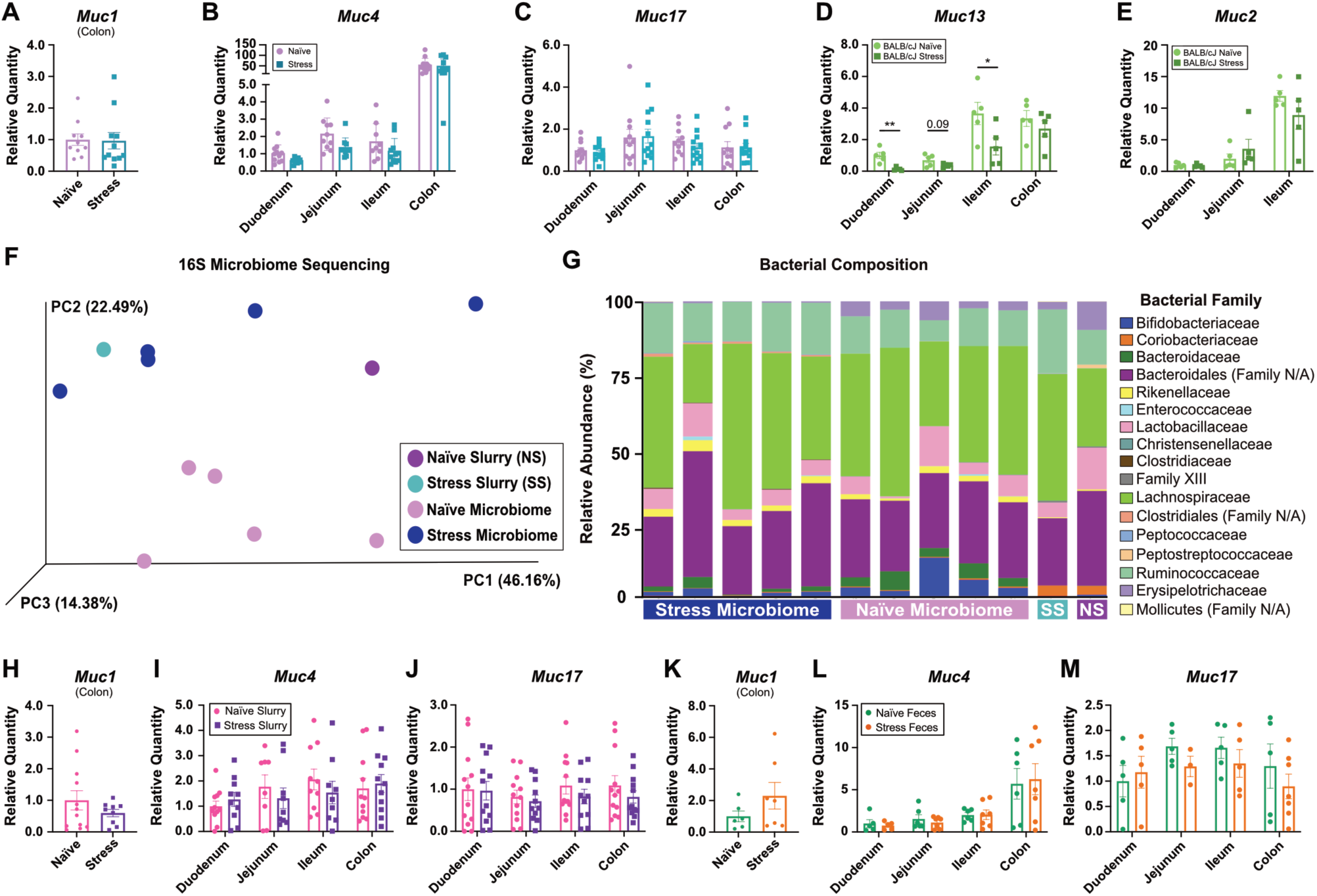
Mucin Expression after Stress and Microbiome Transfers: (A) Relative quantity of *Muc1* expression in colon of naïve and stress mice by qPCR. *Muc*1 was not detected in other sections (two tailed t-test). Relative quantities of (B) *Muc4* and (C) *Muc17* expression in the intestines of naïve and stress mice by qPCR. Two-way ANOVA (D) Relative quantity of *Muc13* and (E) *Muc2* expression in sections of the intestines from naïve and stressed male BALB/cJ mice by qPCR. Two-way ANOVA. Male mice. Experiments A-C, n=11/12 per group. Experiments D-E, n=5 per group. N=1. (F) 16S PCoA plot between naïve and stress slurries and naïve and stress microbiome recipients. (G) Individual bacterial family compositions. Male mice, n=5/group, slurry samples from gavage microbiome mixture, N=1. Relative quantities of (H) *Muc1*, (I) *Muc4*, and (J) *Muc17* from the intestines of mice treated with antibiotics then given naïve or stress fecal slurry. Male mice, 12/group, N=1, Two-way ANOVA. Relative quantities of (K) *Muc1*, (L) *Muc4*, and (M) *Muc17* in germ free mice given naïve or stress feces. Male mice, 7/group, N=1, Two-way ANOVA.

**Extended Data 3:**
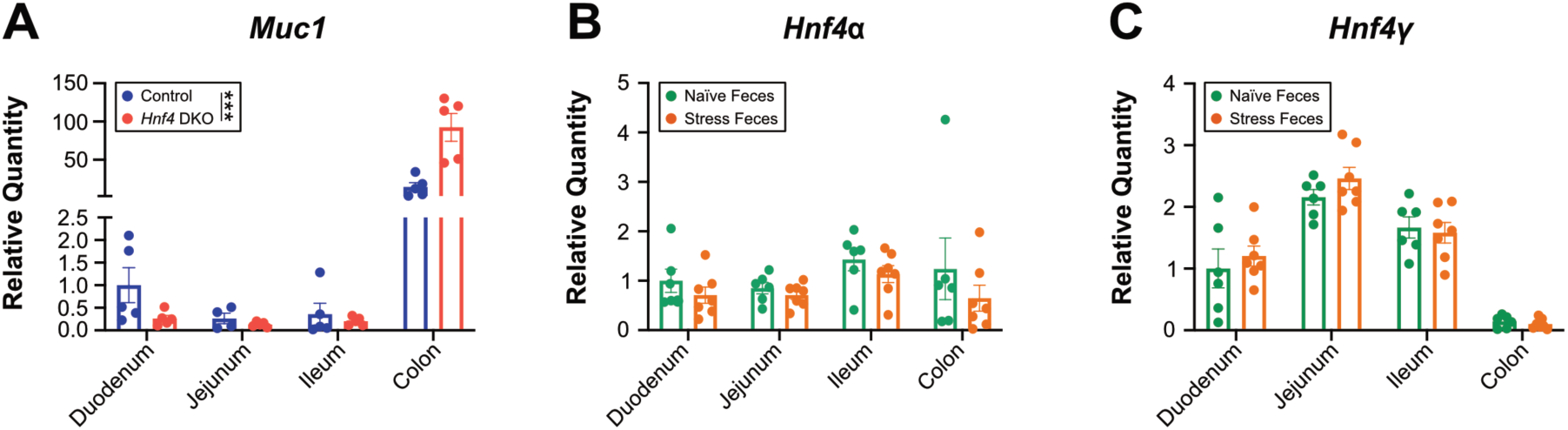
HNF4 Regulates Mucins Independently of Transferrable Microbial Products. Relative quantities of (A) *Muc1* in the intestines of *Hnf4* DKO mice. Female mice, 5/group. Two-way ANOVA. Relative quantities of (B) *Hnf4α* and (C) *Hnf4ψ* in the intestines of germ-free mice given stress or naïve feces. Male mice, 7/group. Two-way ANOVA.

**Extended Data 4:**
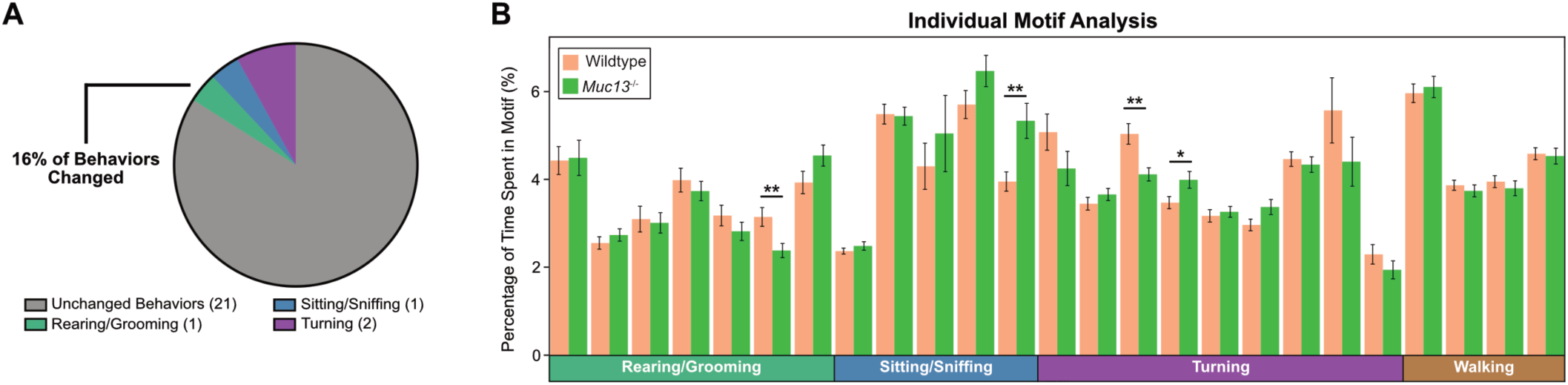
DeepLabCut Analysis of Baseline Behaviors: (A) Pie chart representing quantified behavioral motifs changed between wildtype and *Muc13*^-/-^ animals. (B) Individual motif analysis comparing baseline behaviors in wildtype and *Muc13*^-/-^ animals. Two tailed t-tests, male and female mice, n=13-24 per group, representative of 3 experiments.

**Extended Data 5: Data Tables:** (1-MS Transitions): Table of mass over charge number (m/z) for analytes examined between naïve and stressed animals. (2-NaivevStress_Genus_WilcoxRank): Table of naïve vs stress genus level Wilcox Rank Sum statistics (for Ext. Data 1J). (3-ASV+Genus_ControltoMuc13KO): Table of statistics for ASV and Genus level changes between baseline control, baseline *Muc13^-/-^*, 1wk UCMRS control, 1wk UCMRS *Muc13^-/-^* animals. (4-qPCR Probes): Table of qPCR probes used in manuscript. (5-Statistics Table): Table of statistics for all data in manuscript.

## Notes

### Competing Interest Statement

The authors have declared no competing interest.

### Summary of Updates

The first name of one of the authors was misspelled.

## References

1 Santomauro, D. F. et al. Global prevalence and burden of depressive and anxiety disorders in 204 countries and territories in 2020 due to the COVID-19 pandemic. The Lancet 398, 1700–1712, doi:https://doi.org/10.1016/S0140-6736(21)02143-7 (2021).

2 Fava, M. Diagnosis and definition of treatment-resistant depression. Biological Psychiatry 53, 649–659, doi:https://doi.org/10.1016/S0006-3223(03)00231-2 (2003).

3 Heim, C. & Binder, E. B. Current research trends in early life stress and depression: Review of human studies on sensitive periods, gene–environment interactions, and epigenetics. Experimental Neurology 233, 102–111, doi:https://doi.org/10.1016/j.expneurol.2011.10.032 (2012).

4 McGonagle, K. A. & Kessler, R. C. Chronic stress, acute stress, and depressive symptoms. American Journal of Community Psychology 18, 681–706, doi:https://doi.org/10.1007/BF00931237 (1990).

5 Lach, G., Schellekens, H., Dinan, T. G. & Cryan, J. F. Anxiety, Depression, and the Microbiome: A Role for Gut Peptides. Neurotherapeutics : the journal of the American Society for Experimental NeuroTherapeutics 15, 36–59, doi:10.1007/s13311-017-0585-0 (2018).

6 Foster, J. A. & McVey Neufeld, K. A. Gut-brain axis: how the microbiome influences anxiety and depression. Trends in neurosciences 36, 305–312, doi:10.1016/j.tins.2013.01.005 (2013).

7 Marin, I. A. et al. Microbiota alteration is associated with the development of stress-induced despair behavior. Scientific reports 7, 43859–43859, doi:10.1038/srep43859 (2017).

8 Tian, P. et al. Bifidobacterium breve CCFM1025 attenuates major depression disorder via regulating gut microbiome and tryptophan metabolism: A randomized clinical trial. Brain, behavior, and immunity 100, 233–241, doi:https://doi.org/10.1016/j.bbi.2021.11.023 (2022).

9 Berding, K. & Cryan, J. F. Microbiota-targeted interventions for mental health. Curr Opin Psychiatry 35, 3–9, doi:10.1097/yco.0000000000000758 (2022).

10 Valles-Colomer, M. et al. The neuroactive potential of the human gut microbiota in quality of life and depression. Nature Microbiology 4, 623–632, doi:10.1038/s41564-018-0337-x (2019).

11 Park, S.-Y. et al. Strain-level fitness in the gut microbiome is an emergent property of glycans and a single metabolite. Cell 185, 513–529.e521, doi:10.1016/j.cell.2022.01.002 (2022).

12 Zmora, N. et al. Personalized Gut Mucosal Colonization Resistance to Empiric Probiotics Is Associated with Unique Host and Microbiome Features. Cell 174, 1388–1405.e1321, doi:https://doi.org/10.1016/j.cell.2018.08.041 (2018).

13 Borisova, M. A. et al. Mucin-2 knockout is a model of intercellular junction defects, mitochondrial damage and ATP depletion in the intestinal epithelium. Scientific reports 10, 21135, doi:10.1038/s41598-020-78141-4 (2020).

14 Hansson, G. C. Mucins and the Microbiome. Annual Review of Biochemistry 89, 769–793, doi:10.1146/annurev-biochem-011520-105053 (2020).

15 Johansson, M. E. V. et al. Composition and functional role of the mucus layers in the intestine. Cellular and Molecular Life Sciences 68, 3635, doi:10.1007/s00018-011-0822-3 (2011).

16 Van der Sluis, M. et al. Muc2-Deficient Mice Spontaneously Develop Colitis, Indicating That MUC2 Is Critical for Colonic Protection. Gastroenterology 131, 117–129, doi:https://doi.org/10.1053/j.gastro.2006.04.020 (2006).

17 Velcich, A. et al. Colorectal cancer in mice genetically deficient in the mucin Muc2. Science (New York, N.Y.) 295, 1726–1729, doi:10.1126/science.1069094 (2002).

18 Johansson, M. E. V. a. H. G. C. The Mucins Vol. 2 381–388 (2016).

19 Pelaseyed, T. & Hansson, G. C. Membrane mucins of the intestine at a glance. Journal of Cell Science 133, jcs240929, doi:10.1242/jcs.240929 (2020).

20 Van Herreweghen, F., De Paepe, K., Marzorati, M. & Van de Wiele, T. Mucin as a Functional Niche is a More Important Driver of in Vitro Gut Microbiota Composition and Functionality than Supplementation of Akkermansia m uciniphila. Applied and environmental microbiology 87, doi:10.1128/aem.02647-20 (2020).

21 Duncan, K., Carey-Ewend, K. & Vaishnava, S. Spatial analysis of gut microbiome reveals a distinct ecological niche associated with the mucus layer. Gut Microbes, 1–21, doi:10.1080/19490976.2021.1874815 (2021).

22 Sicard, J.-F., Le Bihan, G., Vogeleer, P., Jacques, M. & Harel, J. Interactions of Intestinal Bacteria with Components of the Intestinal Mucus. Frontiers in cellular and infection microbiology 7, 387–387, doi:10.3389/fcimb.2017.00387 (2017).

23 Werlang, C., Cárcarmo-Oyarce, G. & Ribbeck, K. Engineering mucus to study and influence the microbiome. Nature Reviews Materials 4, 134–145, doi:10.1038/s41578-018-0079-7 (2019).

24 Sommer, F. et al. Altered Mucus Glycosylation in Core 1 O-Glycan-Deficient Mice Affects Microbiota Composition and Intestinal Architecture. PloS one 9, e85254, doi:10.1371/journal.pone.0085254 (2014).

25 Fu, J. et al. Loss of intestinal core 1-derived O-glycans causes spontaneous colitis in mice. J Clin Invest 121, 1657–1666, doi:10.1172/jci45538 (2011).

26 Biol-N’garagba, M.-C., Niepceron, E., Mathian, B. & Louisot, P. Glucocorticoid-induced maturation of glycoprotein galactosylation and fucosylation processes in the rat small intestine. The Journal of Steroid Biochemistry and Molecular Biology 84, 411–422, doi:https://doi.org/10.1016/S0960-0760(03)00062-1 (2003).

27 Silva, S. D. et al. Stress disrupts intestinal mucus barrier in rats via mucin O-glycosylation shift: prevention by a probiotic treatment. American Journal of Physiology-Gastrointestinal and Liver Physiology 307, G420–G429, doi:10.1152/ajpgi.00290.2013 (2014).

28 Gong, S. et al. Dynamics and correlation of serum cortisol and corticosterone under different physiological or stressful conditions in mice. PloS one 10, e0117503–e0117503, doi:10.1371/journal.pone.0117503 (2015).

29 Luo, Y. et al. Gut microbiota regulates mouse behaviors through glucocorticoid receptor pathway genes in the hippocampus. Translational psychiatry 8, 187, doi:10.1038/s41398-018-0240-5 (2018).

30 Dodiya, H. B. et al. Chronic stress-induced gut dysfunction exacerbates Parkinson’s disease phenotype and pathology in a rotenone-induced mouse model of Parkinson’s disease. Neurobiology of disease 135, 104352, doi:10.1016/j.nbd.2018.12.012 (2020).

31 Saldanha, D., Kumar, N., Ryali, V., Srivastava, K. & Pawar, A. A. Serum Serotonin Abnormality in Depression. Medical Journal Armed Forces India 65, 108–112, doi:https://doi.org/10.1016/S0377-1237(09)80120-2 (2009).

32 Pfleiderer, B. et al. Effective electroconvulsive therapy reverses glutamate/glutamine deficit in the left anterior cingulum of unipolar depressed patients. Psychiatry Research: Neuroimaging 122, 185–192, doi:https://doi.org/10.1016/S0925-4927(03)00003-9 (2003).

33 Bailey, M. T. et al. Exposure to a social stressor alters the structure of the intestinal microbiota: implications for stressor-induced immunomodulation. Brain, behavior, and immunity 25, 397–407, doi:10.1016/j.bbi.2010.10.023 (2011).

34 Bastiaanssen, T. F. S. et al. Gutted! Unraveling the Role of the Microbiome in Major Depressive Disorder. Harv Rev Psychiatry 28, 26–39, doi:10.1097/HRP.0000000000000243 (2020).

35 Li, N. et al. Fecal microbiota transplantation from chronic unpredictable mild stress mice donors affects anxiety-like and depression-like behavior in recipient mice via the gut microbiota-inflammation-brain axis. Stress 22, 592–602, doi:10.1080/10253890.2019.1617267 (2019).

36 Karl, J. P. et al. Changes in intestinal microbiota composition and metabolism coincide with increased intestinal permeability in young adults under prolonged physiological stress. American journal of physiology. Gastrointestinal and liver physiology 312, G559–g571, doi:10.1152/ajpgi.00066.2017 (2017).

37 Hansson, G. C. Mucins and the Microbiome. Annu Rev Biochem, doi:10.1146/annurev-biochem-011520-105053 (2020).

38 Sicard, J. F., Le Bihan, G., Vogeleer, P., Jacques, M. & Harel, J. Interactions of Intestinal Bacteria with Components of the Intestinal Mucus. Frontiers in cellular and infection microbiology 7, 387, doi:10.3389/fcimb.2017.00387 (2017).

39 Merchak, A. R. et al. Lactobacillus maintains IFNγ homeostasis to promote behavioral stress resilience. bioRxiv, 2023.2005.2010.540223, doi:10.1101/2023.05.10.540223 (2023).

40 Johansson, M. E. et al. Normalization of Host Intestinal Mucus Layers Requires Long-Term Microbial Colonization. Cell Host Microbe 18, 582–592, doi:10.1016/j.chom.2015.10.007 (2015).

41 Kent, W. J. et al. The human genome browser at UCSC. Genome Res 12, 996–1006, doi:10.1101/gr.229102 (2002).

42 Chahar, S. et al. Chromatin profiling reveals regulatory network shifts and a protective role for hepatocyte nuclear factor 4α during colitis. Mol Cell Biol 34, 3291–3304, doi:10.1128/mcb.00349-14 (2014).

43 Lickwar, C. R. et al. Transcriptional Integration of Distinct Microbial and Nutritional Signals by the Small Intestinal Epithelium. Cellular and molecular gastroenterology and hepatology 14, 465–493, doi:https://doi.org/10.1016/j.jcmgh.2022.04.013 (2022).

44 Babeu, J. P. & Boudreau, F. Hepatocyte nuclear factor 4-alpha involvement in liver and intestinal inflammatory networks. World journal of gastroenterology 20, 22–30, doi:10.3748/wjg.v20.i1.22 (2014).

45 Chen, L. et al. The nuclear receptor HNF4 drives a brush border gene program conserved across murine intestine, kidney, and embryonic yolk sac. Nature Communications 12, 2886, doi:10.1038/s41467-021-22761-5 (2021).

46 Shurer, C. R. et al. Physical Principles of Membrane Shape Regulation by the Glycocalyx. Cell 177, 1757–1770.e1721, doi:https://doi.org/10.1016/j.cell.2019.04.017 (2019).

47 Lu, H. et al. Crosstalk of hepatocyte nuclear factor 4a and glucocorticoid receptor in the regulation of lipid metabolism in mice fed a high-fat-high-sugar diet. Lipids in Health and Disease 21, 46, doi:10.1186/s12944-022-01654-6 (2022).

48 Chen, L. et al. A reinforcing HNF4-SMAD4 feed-forward module stabilizes enterocyte identity. Nat Genet 51, 777–785, doi:10.1038/s41588-019-0384-0 (2019).

49 Chen, L. et al. Three-dimensional interactions between enhancers and promoters during intestinal differentiation depend upon HNF4. Cell reports 34, doi:10.1016/j.celrep.2020.108679 (2021).

50 Chen, L. et al. HNF4 Regulates Fatty Acid Oxidation and Is Required for Renewal of Intestinal Stem Cells in Mice. Gastroenterology 158, 985–999.e989, doi:10.1053/j.gastro.2019.11.031 (2020).

51 Gupta, B. K. et al. Functions and regulation of MUC13 mucin in colon cancer cells. J Gastroenterol 49, 1378–1391, doi:10.1007/s00535-013-0885-z (2014).

52 Darsigny, M. et al. Hepatocyte Nuclear Factor-4α Promotes Gut Neoplasia in Mice and Protects against the Production of Reactive Oxygen Species. Cancer research 70, 9423–9433, doi:10.1158/0008-5472.Can-10-1697 (2010).

53 Kiselyuk, A. et al. HNF4α antagonists discovered by a high-throughput screen for modulators of the human insulin promoter. Chem Biol 19, 806–818, doi:10.1016/j.chembiol.2012.05.014 (2012).

54 Lee, S. H., Veeriah, V. & Levine, F. A potent HNF4α agonist reveals that HNF4α controls genes important in inflammatory bowel disease and Paneth cells. PloS one 17, e0266066, doi:10.1371/journal.pone.0266066 (2022).

55 Gurumurthy, C. B. et al. Creation of CRISPR-based germline-genome-engineered mice without ex vivo handling of zygotes by i-GONAD. Nature Protocols 14, 2452–2482, doi:10.1038/s41596-019-0187-x (2019).

56 Ohtsuka, M. et al. i-GONAD: a robust method for in situ germline genome engineering using CRISPR nucleases. Genome Biology 19, 25, doi:10.1186/s13059-018-1400-x (2018).

57 Malaker, S. A. et al. The mucin-selective protease StcE enables molecular and functional analysis of human cancer-associated mucins. Proceedings of the National Academy of Sciences 116, 7278–7287, doi:10.1073/pnas.1813020116 (2019).

58 Malaker, S. A. et al. Revealing the human mucinome. Nature Communications 13, 3542, doi:10.1038/s41467-022-31062-4 (2022).

59 Pothion, S., Bizot, J.-C., Trovero, F. & Belzung, C. Strain differences in sucrose preference and in the consequences of unpredictable chronic mild stress. Behavioural brain research 155, 135–146, doi:https://doi.org/10.1016/j.bbr.2004.04.008 (2004).

60 Peng, Y.-L. et al. Inducible nitric oxide synthase is involved in the modulation of depressive behaviors induced by unpredictable chronic mild stress. Journal of neuroinflammation 9, 75, doi:10.1186/1742-2094-9-75 (2012).

61 Nollet, M., Guisquet, A.-M. L. & Belzung, C. Models of Depression: Unpredictable Chronic Mild Stress in Mice. Current protocols in pharmacology 61, 5.65.61–65.65.17, doi:https://doi.org/10.1002/0471141755.ph0565s61 (2013).

62 Rivet-Noor, C. R. et al. Stress-induced despair behavior develops independently of the Ahr-RORγt axis in CD4 + cells. Scientific reports 12, 8594, doi:10.1038/s41598-022-12464-2 (2022).

63 Mathis, A. et al. DeepLabCut: markerless pose estimation of user-defined body parts with deep learning. Nature Neuroscience 21, 1281–1289, doi:10.1038/s41593-018-0209-y (2018).

64 Bergstrom, K. et al. Proximal colon–derived O-glycosylated mucus encapsulates and modulates the microbiota. Science (New York, N.Y.) 370, 467–472, doi:10.1126/science.aay7367 (2020).

65 Wu, M. et al. The Dynamic Changes of Gut Microbiota in Muc2 Deficient Mice. International journal of molecular sciences 19, doi:10.3390/ijms19092809 (2018).

66 Bergstrom, K. S. et al. Muc2 protects against lethal infectious colitis by disassociating pathogenic and commensal bacteria from the colonic mucosa. PLoS pathogens 6, e1000902, doi:10.1371/journal.ppat.1000902 (2010).

67 Johansson, M. E., Thomsson, K. A. & Hansson, G. C. Proteomic analyses of the two mucus layers of the colon barrier reveal that their main component, the Muc2 mucin, is strongly bound to the Fcgbp protein. J Proteome Res 8, 3549–3557, doi:10.1021/pr9002504 (2009).

68 Johansson, M. E. V. et al. The inner of the two Muc2 mucin-dependent mucus layers in colon is devoid of bacteria. Proceedings of the National Academy of Sciences 105, 15064–15069, doi:10.1073/pnas.0803124105 (2008).

69 Marczynski, M. et al. Structural Alterations of Mucins Are Associated with Losses in Functionality. Biomacromolecules, doi:10.1021/acs.biomac.1c00073 (2021).

70 Ware, E. B. et al. Comparative genome-wide association studies of a depressive symptom phenotype in a repeated measures setting by race/ethnicity in the multi-ethnic study of atherosclerosis. BMC Genetics 16, 118, doi:10.1186/s12863-015-0274-0 (2015).

71 Dunn, E. C. et al. Genome-wide association study of depressive symptoms in the Hispanic Community Health Study/Study of Latinos. Journal of psychiatric research 99, 167–176, doi:10.1016/j.jpsychires.2017.12.010 (2018).

72 Rivet-Noor, C. & Gaultier, A. The Role of Gut Mucins in the Etiology of Depression. Front Behav Neurosci 14, 592388, doi:10.3389/fnbeh.2020.592388 (2020).

73 McClain, L. L. et al. Rare variants and biological pathways identified in treatment-refractory depression. J Neurosci Res 98, 1322–1334, doi:10.1002/jnr.24609 (2020).

74 Vető, B. et al. The transcriptional activity of hepatocyte nuclear factor 4 alpha is inhibited via phosphorylation by ERK1/2. PloS one 12, e0172020, doi:10.1371/journal.pone.0172020 (2017).

75 Gourley, S. L., Wu, F. J. & Taylor, J. R. Corticosterone Regulates pERK1/2 Map Kinase in a Chronic Depression Model. Annals of the New York Academy of Sciences 1148, 509–514, doi:https://doi.org/10.1196/annals.1410.076 (2008).

76 Yang, C.-H., Huang, C.-C. & Hsu, K.-S. Behavioral Stress Modifies Hippocampal Synaptic Plasticity through Corticosterone-Induced Sustained Extracellular Signal-Regulated Kinase/Mitogen-Activated Protein Kinase Activation. The Journal of Neuroscience 24, 11029–11034, doi:10.1523/jneurosci.3968-04.2004 (2004).

77 Yu, J. et al. Involvement of ERK1/2 signalling and growth-related molecules’ expression in response to heat stress-induced damage in rat jejunum and IEC-6 cells. International Journal of Hyperthermia 26, 538–555, doi:10.3109/02656736.2010.481276 (2010).

78 Herath, M., Hosie, S., Bornstein, J. C., Franks, A. E. & Hill-Yardin, E. L. The Role of the Gastrointestinal Mucus System in Intestinal Homeostasis: Implications for Neurological Disorders. Frontiers in cellular and infection microbiology 10, doi:10.3389/fcimb.2020.00248 (2020).

79 Tyanova, S., Temu, T. & Cox, J. The MaxQuant computational platform for mass spectrometry-based shotgun proteomics. Nature Protocols 11, 2301–2319, doi:10.1038/nprot.2016.136 (2016).

80 Kozich, J. J., Westcott, S. L., Baxter, N. T., Highlander, S. K. & Schloss, P. D. Development of a dual-index sequencing strategy and curation pipeline for analyzing amplicon sequence data on the MiSeq Illumina sequencing platform. Applied and environmental microbiology 79, 5112–5120, doi:10.1128/aem.01043-13 (2013).

81 Team, R. C. R: A language and environment for statistical computing. R Foundation for Statistical Computing. doi:https://www.R-project.org/. (2021).

82 Callahan, B. J. et al. DADA2: High-resolution sample inference from Illumina amplicon data. Nat Methods 13, 581–583, doi:10.1038/nmeth.3869 (2016).

83 Quast, C. et al. The SILVA ribosomal RNA gene database project: improved data processing and web-based tools. Nucleic Acids Res 41, D590–596, doi:10.1093/nar/gks1219 (2013).

84 Jari Oksanen, F. G. B., Michael Friendly, Roeland Kindt, Pierre Legendre, Dan McGlinn, Peter R. & Minchin, R. B. O. H., Gavin L. Simpson, Peter Solymos, M. Henry H. Stevens, Eduard Szoecs and Helene Wagner. vegan: Community Ecology Package. R package version 2.5-7. doi:https://CRAN.R-project.org/package=vegan (2020).

85 McMurdie, P. J. & Holmes, S. phyloseq: An R Package for Reproducible Interactive Analysis and Graphics of Microbiome Census Data. PloS one 8, e61217, doi:10.1371/journal.pone.0061217 (2013).

86 Leo Lahti, F. G. M. E., Sudarshan Shetty, Tuomas Borman, Domenick J. Braccia, Ruizhu Huang, Hector Corrada Bravo. Microbiome R package. doi:http://microbiome.github.io.

87 Hadley Wickham, M. A., Jennifer Bryan, Winston Cheng, Lucy D’Agostino, Romain François, Garrett Grolemund, Alex Hayes, Lionel Henry, Max Kuhn, Thomas Lin Pedersen. Welcome to the Tidyverse. Journal of Open Source Software 4, 1686, doi:https://doi.org/10.21105/joss.01686 (2019).

88 Wickham, H. ggplot2: Elegant Graphics for Data Analysis. (Springer-Verlag 2016).

89 Robin, X. et al. pROC: an open-source package for R and S+ to analyze and compare ROC curves. BMC Bioinformatics 12, 77, doi:10.1186/1471-2105-12-77 (2011).

90 Classification and regression by randomForest (R news 2, 2002).

91 Kuhn, M. caret: Classification and Regression Training. R package version 6.0-90 (2021).

92 Luxem, K., Fuhrmann, F., Kürsch, J., Remy, S. & Bauer, P. Identifying Behavioral Structure from Deep Variational Embeddings of Animal Motion. bioRxiv, 2020.2005.2014.095430, doi:10.1101/2020.05.14.095430 (2020).

93 Concordet, J.-P. & Haeussler, M. CRISPOR: intuitive guide selection for CRISPR/Cas9 genome editing experiments and screens. Nucleic Acids Research 46, W242–W245, doi:10.1093/nar/gky354 (2018).

